# Parvalbumin interneurons and dentate gyrus homeostatic dysregulation shape epileptogenesis in Angelman syndrome model mice

**DOI:** 10.64898/2026.09.17.752419

**Authors:** Nicholas W. Ringelberg, David W. Kipp, Renée E. Mayfield, Lucas M. James, Audrey L. Smith, Paul B. Manis, Alain C. Burette, Michael R. Kasten, Benjamin D. Philpot

## Abstract

Understanding how neural circuits transition from seizure-resistant to seizure-prone is essential for developing improved epilepsy therapies. Here, we study this process by leveraging the heightened susceptibility to seizure kindling of Angelman syndrome (AS) model mice, which lack the maternal *Ube3a* (*mUbe3a*) allele. We identify parvalbumin-expressing (PV+) interneurons as critical gatekeepers; selective *mUbe3a* deletion in PV+ neurons phenocopies enhanced AS epileptogenesis, whereas restoring UBE3A broadly in GABAergic neurons confers seizure resistance. Further, pathological remodeling of the extracellular matrix in the dentate gyrus faithfully tracks with post-kindling seizure susceptibility, highlighting this region’s particular relevance to enhanced epileptogenesis. Mechanistically, we uncover a ‘two-hit’ electrophysiologic phenomenon in AS model mice: kindling fails to recruit compensatory inhibition onto dentate granule cells and instead drives their maladaptive intrinsic hyperexcitability. Together, these findings link cell type-specific inhibitory dysfunction and altered homeostatic plasticity to epileptogenesis, suggesting future circuit-based treatment strategies.

## Introduction

Angelman syndrome (AS) is a severe neurodevelopmental disorder affecting approximately 500,000 individuals worldwide (Duis et al., 2022). While life expectancy remains relatively normal, individuals with AS face lifelong challenges, including seizures, motor dysfunction, intellectual disability, and the absence of speech (Buiting et al., 2016; den Besten et al., 2021). Epilepsy in AS is particularly impactful: seizures affect around 90% of AS individuals and are highly refractory to treatment, remaining uncontrolled in up to 77% of patients (Thibert et al., 2009, 2013; Bindels-de Heus et al., 2020). This intractability mirrors the broader global burden of epilepsy, which impacts 50 million people, with 30% experiencing seizures that are refractory to medication despite decades of pharmacological development (Picot et al., 2008; Chen et al., 2018; GBD 2016 Epilepsy Collaborators, 2019; Löscher et al., 2020). A significant barrier to improving outcomes is our limited understanding of epileptogenesis—how neural circuits shift from seizure-resistant to seizure-prone (Goldberg and Coulter, 2013). Consequently, there are currently no preventive therapies to halt epilepsy from developing in high-risk patient populations (Klein and Tyrlikova, 2020; Bebin et al., 2023). The AS mouse model (*Ube3a^m-/p+^*), through its unique seizure profile, offers an ideal opportunity to investigate this gap in knowledge. On the standard C57BL/6J background, this mouse model is characterized by the absence of spontaneous seizures and largely normal seizure thresholds. However, it shows a profound vulnerability to seizure kindling, making it a useful system to dissect the circuit-level mechanisms driving the transition to hyperexcitability (Born et al., 2017; Gu et al., 2019; Egawa et al., 2021).

Here, we investigate the circuit mechanisms that contribute to increased epileptogenesis in AS by manipulating UBE3A expression in specific inhibitory neuron subtypes. We identify a critical role for parvalbumin (PV)-expressing interneurons in preventing epileptogenesis: UBE3A loss in PV+ neurons mimics the increased seizure susceptibility seen in the AS model, while restoring UBE3A across all GABAergic neurons confers strong seizure resistance. Additionally, we demonstrate that enhanced seizure susceptibility is associated with abnormal extracellular matrix remodeling in the hippocampal dentate gyrus. Mechanistically, we find that these circuit changes not only reflect alterations in perisomatic inhibition but also an unexpected increase in the intrinsic excitability of dentate granule cells. These findings suggest a combinatorial role of interneuron dysfunction and altered homeostatic plasticity, leading to circuit hyperexcitability in AS.

## Results

### Loss of UBE3A from parvalbumin interneurons enhances epileptogenesis

The seizure phenotypes of AS model mice are largely driven by loss of UBE3A from inhibitory neurons (Judson et al., 2016; Gu et al., 2019). To determine which inhibitory neuron subtypes influence seizure susceptibility in AS, we generated mice with cell type-specific maternal *Ube3a* deletion from three major forebrain inhibitory populations: neurons expressing parvalbumin (PV), somatostatin (SOM), or vasoactive intestinal peptide (VIP) (Tremblay et al., 2016). We assessed epileptogenesis using the flurothyl kindling paradigm, in which daily generalized seizure inductions, over eight consecutive days, gradually lower myoclonic and generalized seizure thresholds, with the persistent effects of epileptogenesis measured at retest 28 days later (Figure 1A,B) (Samoriski and Applegate, 1997; Kadiyala et al., 2016; Gu et al., 2019).

**Figure 1:**
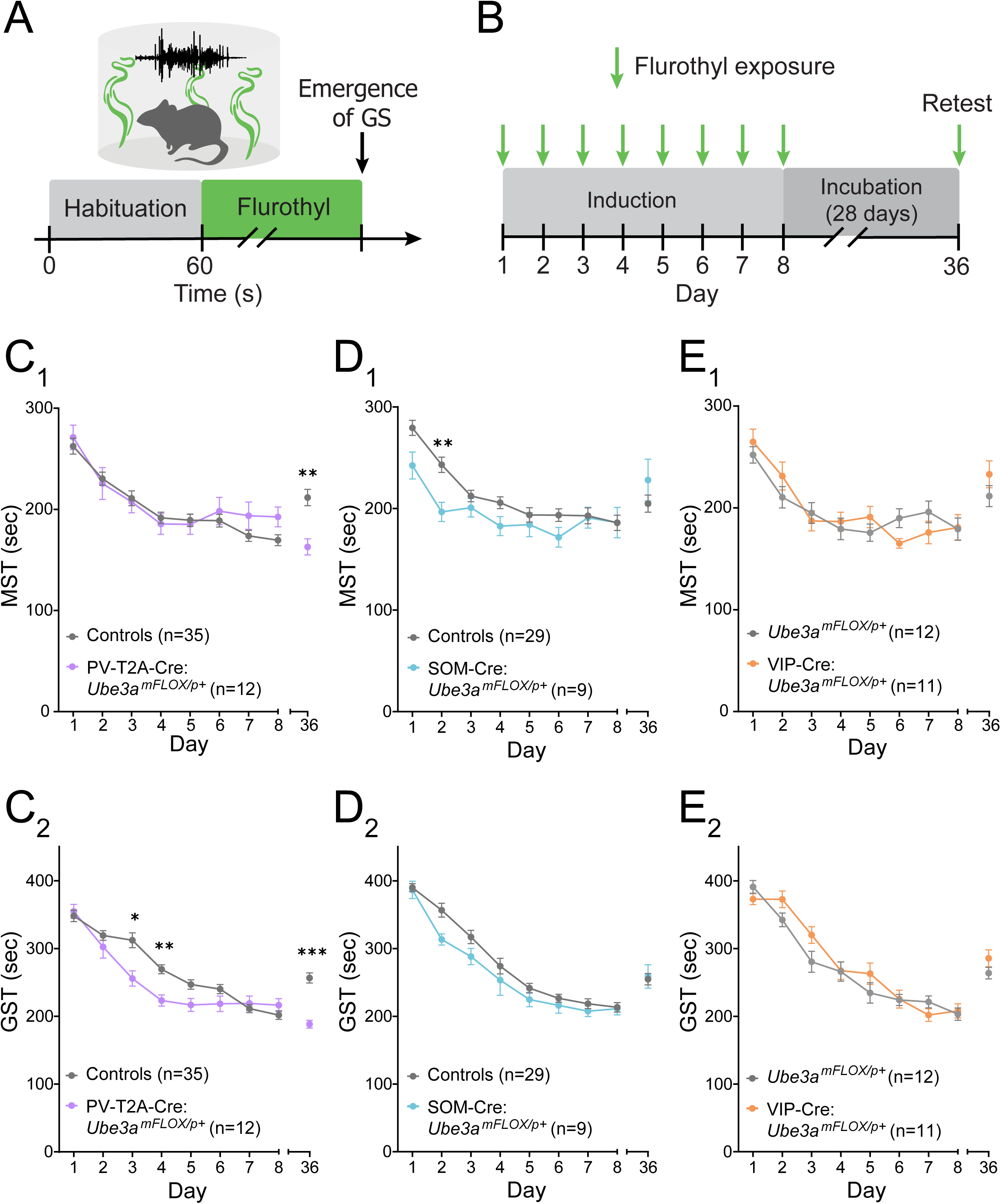
Selective *mUbe3a* deletion from parvalbumin-expressing neurons enhances epileptogenesis. (**A**) Schematic of generalized seizure (GS) induction by flurothyl exposure. (**B**) Schematic of flurothyl kindling timeline. Latency to (**C^1^**) myoclonic (MST) and (**C^2^**) generalized seizure threshold (GST) of PV-T2A-Cre:*Ube3a^mFLOX/p+^* and control mice. (**D^1^,D^2^**) Seizure latencies of SOM-Cre:*Ube3a^mFLOX/p+^* mice. (**E^1^,E^2^**) Seizure latencies of VIP-Cre:*Ube3a^mFLOX/p+^* and control mice. Two-way RM ANOVA with Šídák’s post hoc tests. Data presented as means ± SEM. *P < 0.05, **P < 0.01, ***P<0.001. Control mice for (**C and D**) are WT, *Ube3a^mFLOX/p+^*, and PV-T2A-Cre or SOM-Cre mice. Control groups for (**E**) are *Ube3a^mFLOX/p+^* mice. Control groups are disaggregated in Figure S1.

PV-specific *mUbe3a* deletion, mediated by crossing PV-T2A-Cre mice (Madisen et al., 2012) with conditional *Ube3a^mFLOX^* mice (Judson et al., 2016), robustly decreased seizure threshold on retest and accelerated kindling progression (days 3-4) (Fig. 1C), recapitulating the AS (*Ube3a^m−/p+^*) phenotype. In contrast, *mUbe3a* deletion in SOM+ or VIP+ neurons did not strongly alter kindling progression or retest thresholds (Fig. 1D,E), indicating that PV+ neurons are critical mediators of kindling susceptibility. For these analyses, we made the *a priori* decision to pool data from wildtype (WT), Cre-driver line, and *Ube3a^mFLOX/p+^* mice. However, we note that *Ube3a^mFLOX/p+^* mice exhibited a modest difference in myoclonic seizure threshold on Day 1 of kindling compared to WT mice, despite otherwise showing nearly identical acute seizure latencies and retest thresholds (disaggregated data in Fig. S1A,B). Importantly, PV-T2A-Cre and SOM-Cre mice did not show robust differences in baseline or retest seizure thresholds (Fig. S1C-F).

To validate the role of PV-specific UBE3A loss in driving enhanced epileptogenesis, we replicated this effect of UBE3A loss from PV+ neurons on seizure kindling using an independent PV-Cre driver line (PV-IRES-Cre) (Fig. S2A,B) (Hippenmeyer et al., 2005). Notably, PV-IRES-Cre controls displayed higher seizure thresholds than WT mice upon retest at Day 36, indicating resistance to seizure kindling. Intrigued by this observation, we used immunohistochemistry to find that PV-IRES-Cre mice exhibited ∼40% reduction in hippocampal PV expression with (Fig. S2C,D) or without (Fig. S2E) prior seizure kindling. This effect was replicated with RT-qPCR and western blot of whole brain hemispheres (Fig. S3A,B). We were therefore encouraged to examine PV expression in the PV-T2A-Cre line, where, unexpectedly, we similarly observed this phenomenon on the mRNA and protein levels (Fig. S4A,B). This transcriptional artifact is consistent with reports of decreased endogenous gene expression in other Cre driver lines (Viollet et al., 2017; Joye et al., 2020). Potentially related to its role as a slow-acting calcium buffer, parvalbumin haploinsufficiency has been shown to alter neuronal physiology and animal behavior (Blaustein, 1988; Schwaller et al., 2002, 2004; Eggermann and Jonas, 2011; Wöhr et al., 2015); this phenotype should therefore be considered and appropriately controlled for when interpreting and performing experiments that depend on native calcium dynamics (e.g., imaging) using PV-Cre driver lines. Nonetheless, *mUbe3a* deletion increased kindling susceptibility across both PV driver lines, supporting that loss of UBE3A in this cell type is sufficient to enhance epileptogenic potential.

### PV+ neurons play a major role in GABAergic seizure resilience

Given that pan-GABAergic *mUbe3a* deletion produces severe seizure phenotypes (Judson et al., 2016; Gu et al., 2019), we tested whether GABAergic *mUbe3a* reinstatement could rescue kindling susceptibility in AS mice. We generated mice with pan-GABAergic *mUbe3a* reinstatement by crossing the Gad2-Cre line with the *Ube3a* lox-STOP-lox reinstatement model (*Ube3a^mSTOP^*) (Silva-Santos et al., 2015). First, we confirmed that the *Ube3a^mSTOP/p+^* model mice reproduced the decreased myoclonic and generalized seizure latencies upon retest that have been observed with pan-neuronal UBE3A loss (Fig. 2A,B). In contrast, Gad2-Cre:*Ube3a^mSTOP/p+^* mice displayed remarkable seizure resistance, showing fully rescued seizure latencies on retest and increased thresholds to generalized seizures throughout the kindling process (Fig. 2A,B; disaggregated data in Fig. S5). Furthermore, GABAergic reinstatement protected these mice from seizure-induced death (Fig. 2C), corroborating the high mortality observed in mice with pan-GABAergic deletion of *mUbe3a* (Judson et al., 2016).

**Figure 2:**
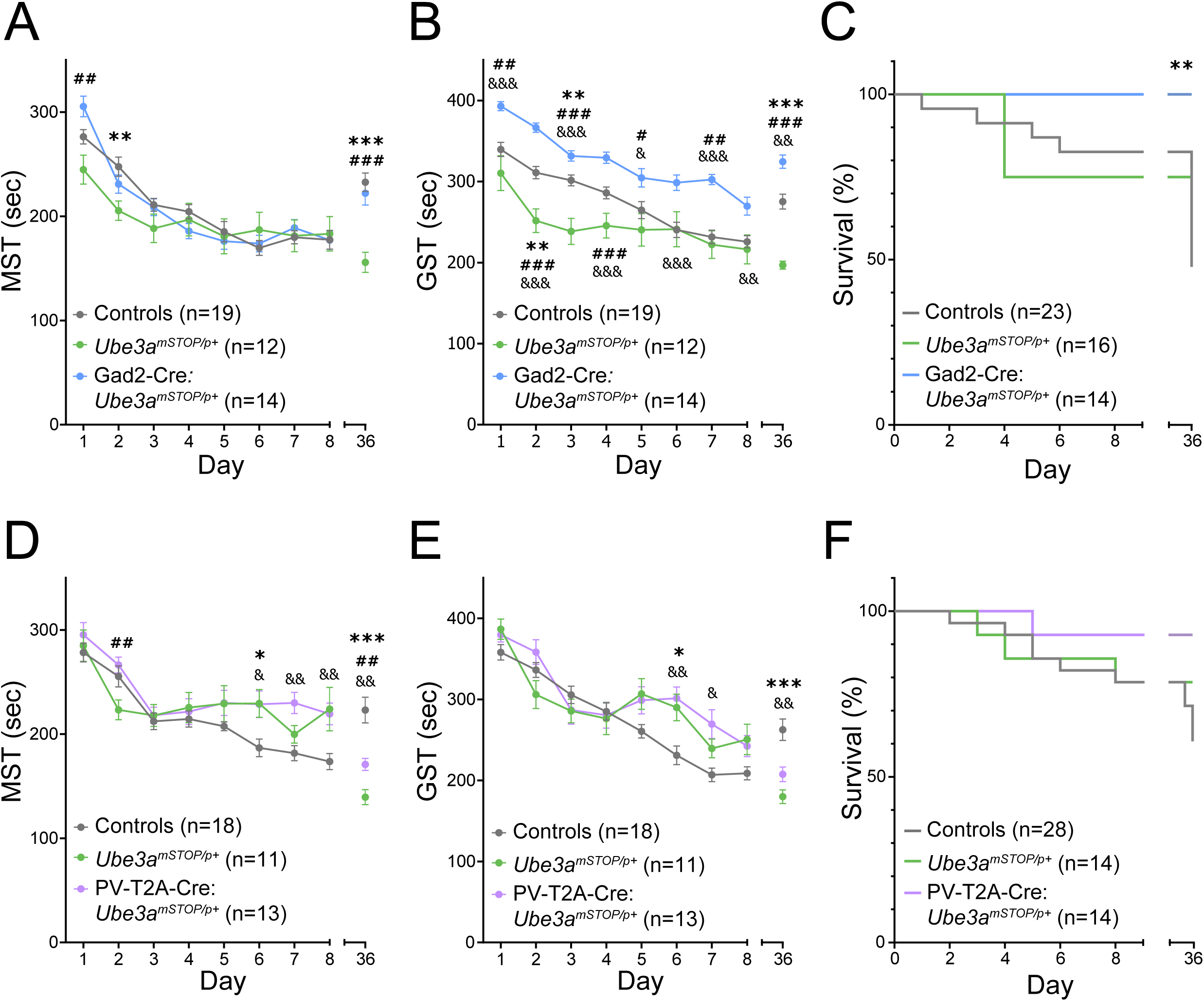
Expression of *mUbe3a* in GABAergic neurons confers resilience to seizure kindling. (**A**) Myoclonic seizure threshold (MST), (**B**) generalized seizure threshold (GST), and (**C**) seizure mortality of Gad2-Cre:*Ube3a^mSTOP/p+^* and control mice. (**D,E**) Seizure latencies and (**F**) mortality of PV-T2A-Cre:*Ube3a^mSTOP/p+^* and control mice. (**A,B,D,E**) Two-way RM ANOVA with Tukey’s post hoc tests. (**C,F**) Log-rank (Mantel-Cox) test. Data presented as means ± SEM. *: Controls vs. *Ube3a^mSTOP/p+^*, #: *Ube3a^mSTOP/p+^* vs. Cre:*Ube3a^mSTOP/p+^*, C: Controls vs. Cre:*Ube3a^mSTOP/p+^*. *P < 0.05, **P < 0.01, ***P<0.001. Control mice are WT and Gad2-Cre or PV-T2A-Cre mice. Control groups are disaggregated in Figure S5.

Next, we tested whether *mUbe3a* reinstatement in PV+ neurons is sufficient to rescue the AS epileptogenic phenotype. To avoid baseline seizure-resistance confounds in the PV-IRES-Cre line, we used PV-T2A-Cre mice for PV+ neuron reinstatement. Reinstatement in PV+ neurons conferred a partial rescue of the myoclonic seizure threshold on retest (Fig. 2D) and a trend toward rescue of the generalized seizure threshold (Fig. 2E). PV-T2A-Cre:*Ube3a^mSTOP/p+^* mice also exhibited a trend toward improved survival during flurothyl kindling, though this did not reach statistical significance (Fig. 2F). Taken together, our results suggest that while UBE3A loss in PV+ neurons plays a major role in seizure susceptibility, loss of UBE3A from additional GABAergic subtypes likely also contributes to kindling susceptibility in AS mice.

### Dentate gyrus extracellular matrix remodeling tracks with seizure susceptibility

The dentate gyrus (DG) functions as a critical gatekeeper limiting hippocampal excitation (Patton and McNaughton, 1995; Amaral et al., 2007; Scharfman, 2019), and DG dysfunction features prominently in epilepsy models (Dengler et al., 2017; Botterill et al., 2019; Lee et al., 2019; Mattis et al., 2022). In AS mice, flurothyl kindling drives a specific pathological signature in the DG: an atypical accumulation of *Wisteria floribunda* agglutinin (WFA)-labeled perineuronal nets (PNNs) in the molecular layer (Gu et al., 2019), a phenomenon later reported in other mouse models of epilepsy (Carstens et al., 2021; Whitebirch et al., 2023; Patel et al., 2024; Woo et al., 2025). To test whether this pathology tracks with seizure susceptibility across genotypes, we quantified WFA fluorescence in our genetic models of seizure susceptibility and resilience.

Consistent with previous findings, kindled *Ube3a^mSTOP/p+^* mice exhibited a robust increase in WFA fluorescence in the DG molecular layer compared to kindled littermate controls (Fig. 3A). Strikingly, pan-GABAergic reinstatement (Gad2-Cre: *Ube3a^mSTOP/p+^*), which conferred complete seizure resilience, also normalized WFA fluorescence intensity to control levels (Fig. 3B). Likewise, *mUbe3a* deletion specifically from PV+ neurons (PV-T2A-Cre), which enhanced seizure susceptibility, was sufficient to drive increased WFA labeling (Fig. 3C). Finally, PV-specific reinstatement resulted in a partial, non-significant, reduction in WFA staining that mirrored the partial seizure behavioral rescue observed in these mice (Fig. 3D). Flurothyl kindled AS mice also show increased astrogliosis in the hippocampus (Judson et al., 2021), a hallmark of seizure-induced pathology (Devinsky et al., 2013). We observed a parallel pattern of reactive astrogliosis, measured by GFAP expression, that tracked with seizure phenotypes across all genotypes, albeit with more modest effect sizes (Fig. S6A–D).

**Figure 3:**
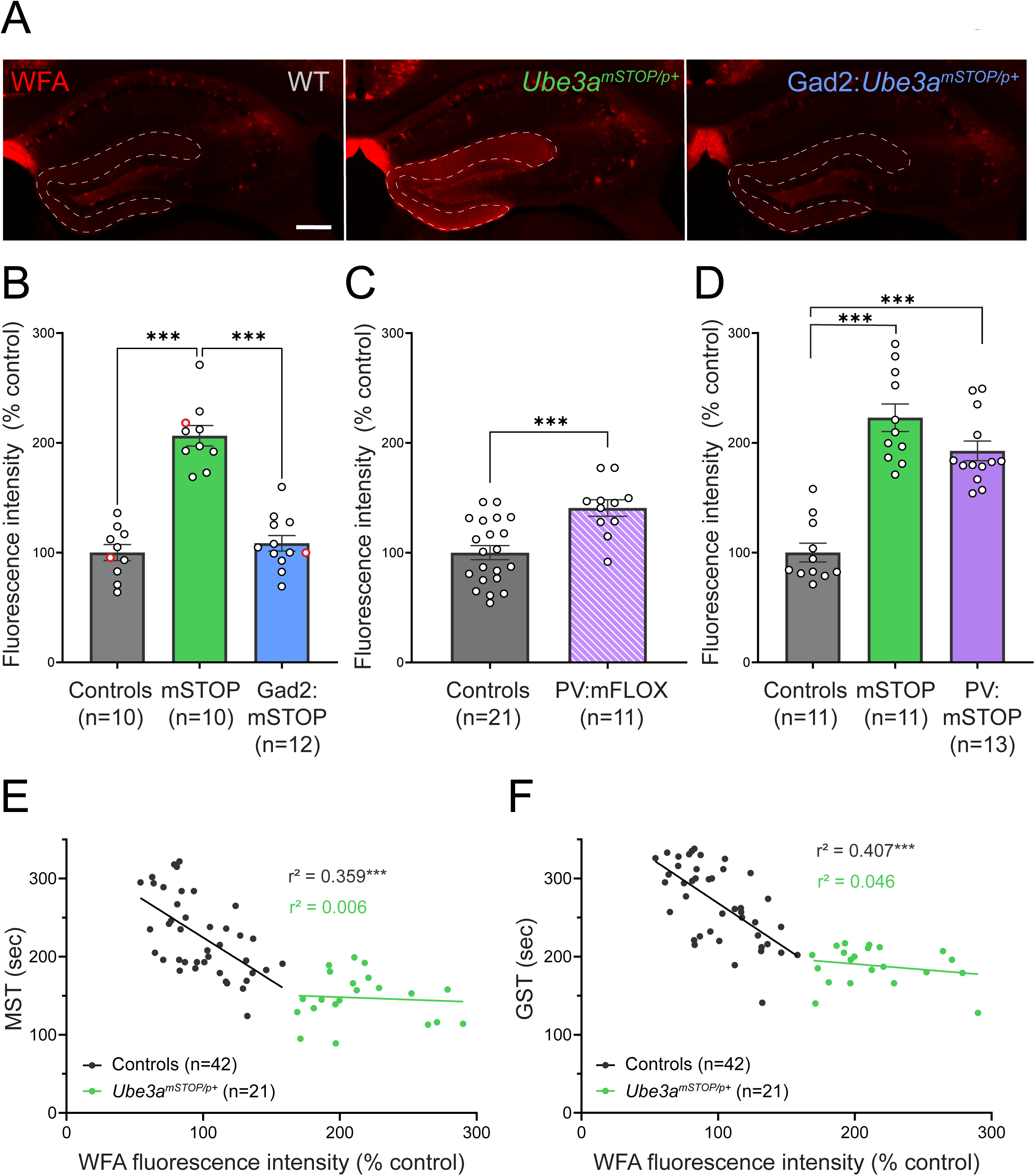
Perineuronal net deposition in the dentate gyrus correlates with seizure susceptibility. (**A**) Representative images of WFA staining in dentate gyrus (DG) of WT, *Ube3a^mSTOP/p+^* mice, and Gad2-Cre:*Ube3a^mSTOP/p+^* mice (Scale bar: 250 µm). (**B**) Mean WFA fluorescence in DG molecular layer, normalized to controls, in flurothyl kindled Gad2-Cre:*Ube3a^mSTOP/p+^* mice, (**C**) PV-T2A-Cre:*Ube3a^mFLOX/p+^* mice, and (**D**) PV-T2A-Cre:*Ube3a^mSTOP/p+^* mice. Red data points in (**B**) correspond to example images in (**A**). (**E**) Correlation analysis of DG WFA fluorescence and myoclonic seizure threshold of controls and *Ube3a^mSTOP/p+^* mice. (**F**) Correlation analysis of DG WFA fluorescence and generalized seizure threshold. (**B,D**) One-way ANOVA with Tukey’s post hoc tests. (**C**) Unpaired two-tailed t-test. (**E,F**) Simple linear regression, with * indicating significantly non-zero slope. Data presented as means ± SEM. *P < 0.05, **P < 0.01, ***P<0.001.

To investigate the relationship between PNNs in the DG and seizure behavior, we correlated WFA fluorescence intensity with retest seizure thresholds of individual mice. In control mice, we observed a strong negative correlation: higher WFA intensity was associated with lower seizure thresholds for both myoclonic (Fig. 3E) and generalized (Fig. 3F) seizures, suggesting an association between PNN accumulation and seizure threshold following kindling. However, WFA intensity did not reliably predict seizure thresholds in *Ube3a^mSTOP/p+^* mice, consistent with a floor effect. A similar negative correlation was found between retest seizure thresholds and GFAP fluorescence intensity (Fig. S6E,F). Overall, these results establish a strong link between seizure-induced pathology in the DG and increased sensitivity to kindling, indicating that the DG is a likely hotspot for conferring enhanced epileptogenic potential.

### Dentate granule cells in AS mice lack the compensatory inhibitory response to seizure kindling

Next, we investigated the physiological consequences of kindling and UBE3A loss in the DG, first examining inhibitory drive onto dentate granule cells. We performed whole-cell patch clamp recordings from dentate granule cells (DGCs) in flurothyl kindled and sham kindled WT and AS mice (Fig. 4A), first assessing differences in spontaneous inhibitory post-synaptic currents (sIPSCs) (Fig. 4B). Surprisingly, sIPSC amplitudes (Fig. 4C), and frequencies (Fig. 4D) were similar between groups, indicating no major difference in basal spontaneous inhibitory synaptic function.

**Figure 4:**
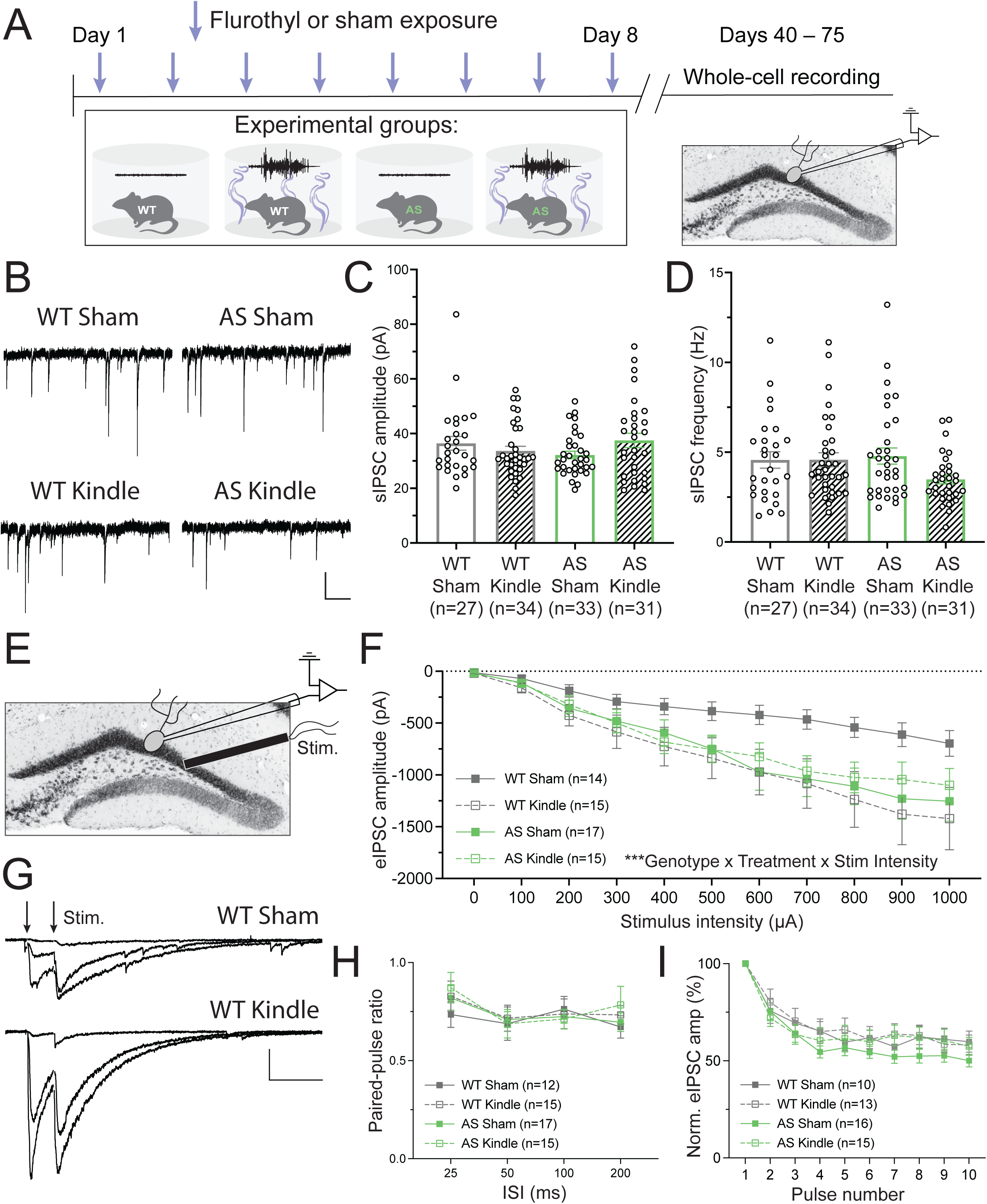
AS mice exhibit occluded perisomatic inhibition onto dentate granule cells following seizure kindling. (**A**) Schematic of flurothyl and sham kindling followed by whole-cell recording. (**B**) Sample traces of sIPSCs in flurothyl and sham kindled WT and AS mice (scale bar: 20 pA, 300 ms). (**C**) Quantification of sIPSC amplitude and (**D**) frequency. Two-way ANOVA. (**E**) Experimental design to record evoked IPSCs onto dentate granule cells. (**F**) Quantification of eIPSC amplitudes. (**G**) Example traces of eIPSCs with 25 ms inter-stimulus interval at stimulation intensities 100, 300, and 700 µA (scale bar: 300 pA, 50 ms). Stimulus artifacts have been removed from example traces. (**H**) Quantification of eIPSC paired-pulse ratio. (**I**) Quantification of eIPSC response to 20 Hz pulse train. (**F,H,I**) Three-way RM ANOVA. Data presented as means ± SEM. n = number of cells. *P < 0.05, **P < 0.01, ***P<0.001.

Given our finding that UBE3A loss in PV+ neurons drives seizure susceptibility, we chose to next focus on perisomatic inhibition, primarily mediated by PV+ basket cells (McBain and Fisahn, 2001; Tremblay et al., 2016). Accordingly, we electrically stimulated the granule cell layer to evoke IPSCs (eIPSCs) from perisomatic targeting interneurons (Fig. 4E,F) and observed a significant three-way interaction. In WT mice, kindling induced an increase in eIPSC amplitude. AS mice, however, displayed increased eIPSC amplitude at baseline, with no further increase in inhibitory strength with kindling, suggesting possible pre-existing synaptic compensation and a resulting ceiling effect. To determine if these inhibitory differences were driven by presynaptic changes, we measured paired-pulse ratios (PPR) across varying inter-stimulus intervals. We detected no significant differences in PPR (Fig. 4H) or in response to 20 Hz pulse trains (Fig. 4I) between groups, suggesting differences in inhibition are likely not driven by changes in release probability. Collectively, these data suggest that AS mice are unable to appropriately increase perisomatic inhibition of DGCs following seizure kindling, either through postsynaptic mechanisms or differences in interneuron excitability.

### Kindling induces hyperexcitability in AS dentate granule cells

We next asked whether kindling alters the intrinsic firing properties of DGCs. Using current-clamp recordings, we found that DGCs from kindled AS mice displayed a robust hyperexcitable phenotype. This was characterized by a leftward shift in the frequency-current (F-I) curve (Fig. 5A), with a corresponding increase in maximum firing gain (Fig. 5B). Importantly, this plasticity was absent in WT mice, indicating genotype-specific sensitivity to kindling. DGCs from kindled AS mice also exhibited other hallmarks of hyperexcitability, including increased input resistance (Fig. 5C) and a trend toward decreased rheobase current (Fig. 5D), with no difference in resting membrane potential, threshold voltage, or maximum firing rate (Table S1). While difficult to directly test, this pattern of hyperexcitability with increased input resistance and similar maximum firing rate is likely explained by the loss of a persistent cation conductance, such as a leak potassium channel conductance or low voltage-activated conductance (Speca et al., 2014; Aggarwal et al., 2021).

**Figure 5:**
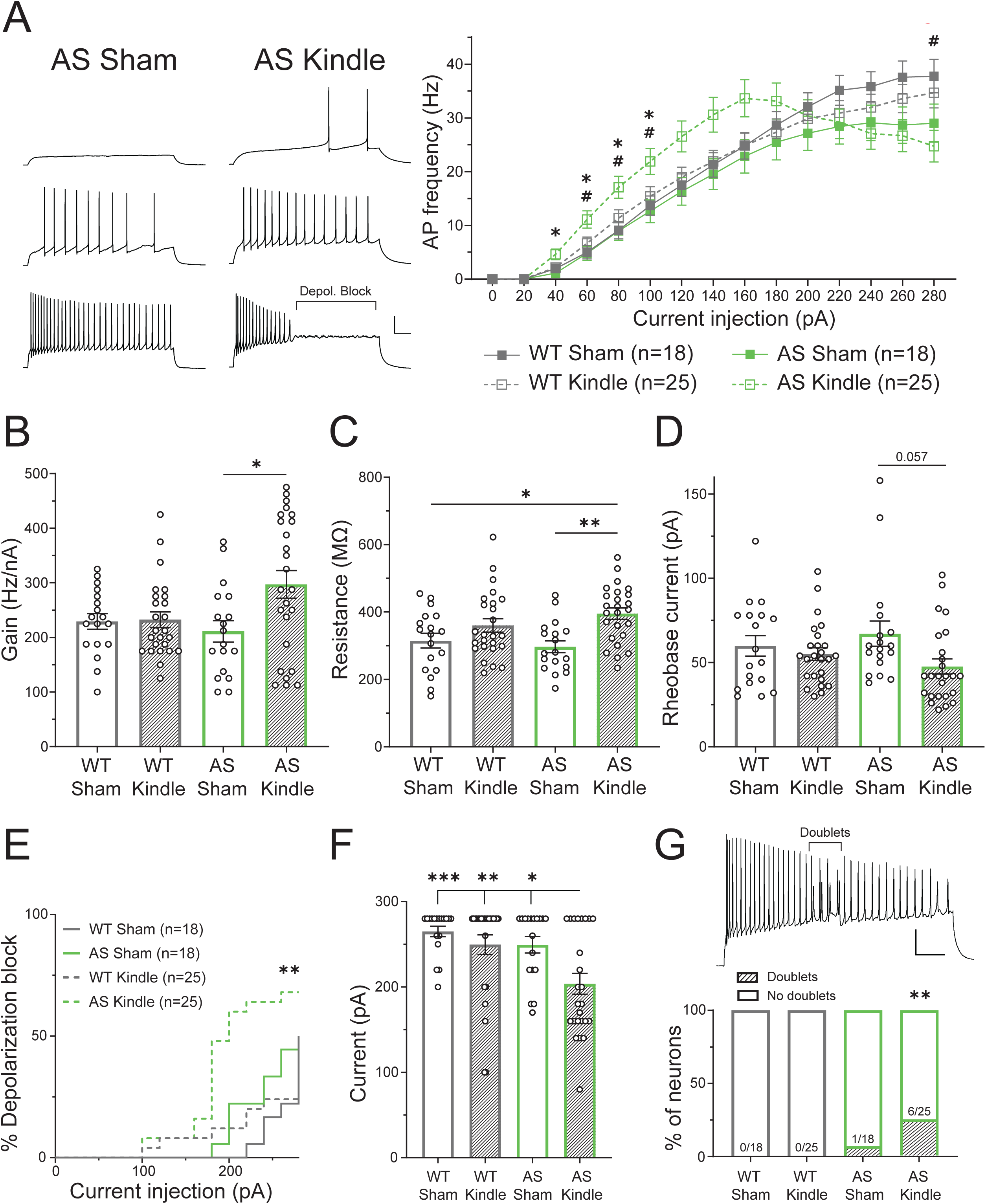
Seizure kindling increases excitability of dentate granule cells of AS mice. (**A**) Current vs firing frequency relationships of flurothyl and sham kindled AS and WT mice.**\** Left: Example current-clamp traces demonstrating increased excitability, and earlier depolarization block in kindled AS mice compared to sham AS. Top: 40 pA, middle: 100 pA, bottom: 240 pA (scale bar: 25 mV, 100 ms). Right: summarized F-I data. Three-way RM ANOVA with Tukey’s post hoc tests. *: AS sham vs AS kindle, #: WT sham vs AS kindle. (**B**) Group comparisons of maximum firing gain, (**C**) input resistance, and (**D**) rheobase current. (**E**) Cumulative incidence of depolarization block across current injections. Log-rank (Mantel-cox) test. (**F**) Quantification of current injection leading to max firing frequency. (**G**) Top: Example current-clamp recording of ‘doublet’ firing in a kindled AS mouse. Bottom: Incidence of observed ‘doublet’ firing across experimental groups. Fisher’s exact test. (**B,C,D,F**) Two-way ANOVA with Tukey’s post hoc tests. Data presented as means ± SEM. n**\** = number of cells. *P < 0.05, **P < 0.01, ***P<0.001.

In addition to increased sensitivity to depolarizing current, DGCs from kindled AS mice displayed an earlier onset of depolarization block (Fig. 5E). Consequently, these cells reached peak firing frequencies at lower current injection steps (Fig. 5F). Notably, a subset of DGCs from kindled AS mice (6 of 25 cells across 5 of 8 mice) exhibited an aberrant ‘doublet’ firing pattern (Fig. 5G). These doublets tended to begin toward the end of the spike train when these cells approached their maximum firing frequencies, and progressed earlier in the spike train with increasing current injections (Fig. S7A-C). In the absence of kindling, doublet firing was exceedingly rare in AS mice; only one DGC from one sham kindled AS mouse (1 of 18 cells from 1 of 6 mice) displayed a doublet firing in response to its highest current injection (Fig. S7D).

To determine whether these doublet-firing cells contribute to the hyperexcitability phenotype of DGCs from kindled AS mice, we separately analyzed their firing properties. Unexpectedly, these doublet firing cells displayed striking hyperexcitability on the F-I curve compared to non-doublet firing cells from kindled AS mice (Fig. 6A). Similarly to the AS kindled neurons when compared with other experimental groups, doublet firing cells displayed a corresponding increase in firing gain (Fig. 6B) and decrease in rheobase current (Fig. 6D). However, these doublet cells also demonstrated firing properties distinct from the global kindled AS group, including increased maximum firing frequency (Fig. 6C), and decreased action potential threshold voltage (Fig. 6E). Doublet firing cells showed comparable input resistance (Fig. 6F), resting membrane potential (Fig. 6G), and spike frequency adaptation (Table S2) to non-doublet cells from this experimental group, and exhibited similar onset of depolarization block (Fig. 6H). This distinct pattern of hyperexcitability, with increased maximum firing frequency but unchanged input resistance, may be driven by changes in ion channels, such as a voltage-gated potassium channel (Kasten et al., 2007; Carver and Shapiro, 2019; Kim et al., 2020). Together, these data indicate that seizure kindling drives pathological hyperexcitability specifically in AS DGCs, likely contributing to their epileptogenic phenotype.

**Figure 6:**
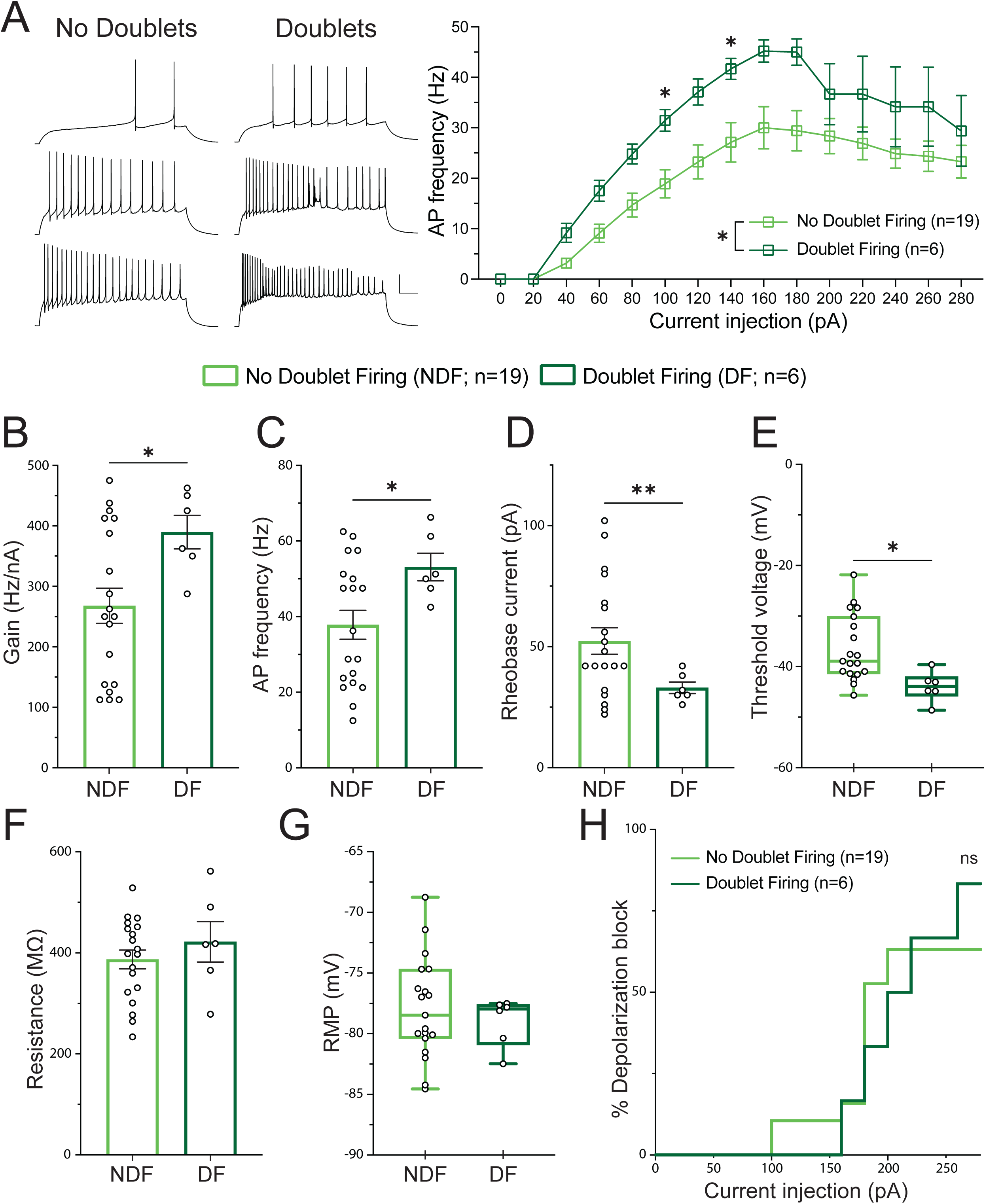
‘Doublet’ firing dentate granule cells demonstrate a distinct hyperexcitability phenotype. (**A**) Current vs firing frequency relationships of doublet and non-doublet firing dentate granule cells from kindled AS mice. Left: Example current-clamp traces demonstrating increased excitability of doublet firing cells. Top: 40 pA, middle: 100 pA, bottom: 160 Pa (scale bar: 25 mV, 100 ms). Right: summarized F-I data. Two-way RM ANOVA with Šídák’s post hoc tests. (**B**) Group comparisons of maximum firing gain, (**C**) maximum firing frequency, (**D**) rheobase current, (**E**) threshold voltage, (**F**) input resistance, and (**G**) resting membrane potential. (**H**) Cumulative incidence of depolarization block across current injections. (**B,C,E,F,G**) Unpaired two-tailed t-test. (**D**) Welch’s t-test. Data presented as means ± SEM. n = number of cells. *P < 0.05, **P < 0.01, ***P<0.001.

## Discussion

Understanding the cell types and circuitry underlying epileptogenesis remains a key challenge in neuroscience with direct clinical importance (Goldberg and Coulter, 2013; Marsh et al., 2021; Simonato et al., 2021; Park, 2024). Here, we leveraged the unique “uncoupled” seizure profile of AS model mice to identify a circuit mechanism that contributes to this transition from seizure-resistant to seizure-prone. Our data reveal a key contribution of PV+ interneurons to this process, serving as crucial gatekeepers to epileptogenesis. PV+ neurons provide critical feedforward perisomatic inhibition via strong excitatory inputs and extensive axonal arbors (Freund and Buzsáki, 1996; Ewell and Jones, 2010; Dengler and Coulter, 2016). This ability to synchronize large populations of principal cells makes them well suited to prevent runaway excitation (Bartos and Elgueta, 2012; Hu et al., 2014; Espinoza et al., 2018), corroborating their demonstrated dysfunction or mislocalization in numerous murine models of epilepsy (Goel et al., 2018; Antoine et al., 2019; Kaneko et al., 2022; Miralles et al., 2024) and human epilepsies (Medici et al., 2016; Wittner and Maglóczky, 2017). Our results demonstrate that perturbation of PV+ neuron development and/or function by loss of UBE3A is sufficient to drive an epileptogenic phenotype without substantially impacting baseline seizure threshold. PV-specific *mUbe3a* reinstatement, however, yielded incomplete rescue relative to pan-GABAergic reinstatement, possibly due to contributions by other interneuron classes, or the relatively delayed expression of parvalbumin, and consequently delayed Cre-mediated *mUbe3a* reinstatement, in the neocortex (de Lecea et al., 1995).

Across multiple genetic manipulations, seizure susceptibility closely correlated with abnormal PNN deposition in the DG molecular layer. While PNNs were first shown to regulate synaptic plasticity in the context of critical window closure in the developing visual cortex (Pizzorusso et al., 2002; Beurdeley et al., 2012), recent studies have reported the emergence of diffuse WFA+ fluorescence in the dentate gyrus in viral encephalitis-mediated epilepsy (Patel et al., 2024), a genetic model of spontaneous recurrent seizures (Carstens et al., 2021), and the pilocarpine model of status epilepticus (Whitebirch et al., 2023; Woo et al., 2025), suggesting a point of convergence on the circuit level. A recent study in one of these models identified DGC hyperexcitability and partially reduced seizure frequency with genetic knockout of aggrecan from DGCs (Patel et al., 2024). Surprisingly, removal of aberrant PNNs in this model did not rescue the DGC hyperexcitability phenotype, suggesting multiple mechanisms are acting synergistically.

Our electrophysiological experiments reveal a dual mechanism potentially underlying increased epileptogenesis in AS mice. First, while kindling in WT mice induced a compensatory increase in perisomatic inhibition onto dentate granule cells, suggesting that neurotypical hippocampal circuitry can thereby maintain stability after seizure induction, this homeostatic adaptation was occluded in AS mice, possibly due to existing circuit compensation. Second, we observed that dentate granule cells from kindled AS mice developed an intrinsic hyperexcitability phenotype, with increased firing gain, higher input resistance, and earlier onset of depolarization block. We also observed the emergence of ‘doublet’ firing in approximately one-quarter of dentate granule cells of kindled AS mice, but these were nearly absent in other groups. To our knowledge, doublet or burst firing patterns have only been observed in DGCs at the beginning of the spike train, where they are posited to facilitate information transfer to CA3 and LTP at the lateral perforant path (Dumenieu et al., 2018; Kim et al., 2023; Shu and Jackson, 2024), suggesting a firing mode that has not been previously characterized. Our observed doublet firing cells were more excitable than non-doublet firing cells, exhibiting increased maximum firing and reduced threshold. The emergence of this subpopulation of particularly hyperexcitable cells raises the possibility that these cells could play an outsized role in local circuit instability, predisposing to seizures. Together, this ‘two-hit’— the occlusion of adaptive perisomatic inhibition combined with the emergence of intrinsic hyperexcitability—provides a circuit-level explanation for the increased susceptibility of AS mice.

While our study elucidates a circuit-level mechanism for epileptogenesis in AS, several questions remain. First, while we establish a key role for PV+ neurons in resilience to epileptogenesis, how UBE3A loss from this population alters their function remains unknown. Second, the causal relationship between PNN accumulation and seizure threshold reduction remains to be established; while the degree of WFA positivity correlates with seizure threshold, direct manipulation of the extracellular matrix will provide important additional insights. Third, while our study focused on the dentate gyrus, due to its striking histopathology in kindled AS mice and its known critical role in regulating seizure propagation (Dengler and Coulter, 2016; Scharfman, 2019), other brain regions likely also contribute to seizure susceptibility in this model.

Despite substantial progress toward new AS treatments, epilepsy in these individuals remains remarkably refractory to current anti-epileptic medications (Thibert et al., 2009; Bindels-de Heus et al., 2020). While multiple UBE3A reinstatement therapies are in clinical development, preclinical studies suggest limited therapeutic windows (Silva-Santos et al., 2015; Milazzo et al., 2021; Rotaru et al., 2023), demonstrating the need for complementary approaches targeting downstream circuit dysfunction. The present study identifies multiple entry points to inform future therapeutic strategies for epilepsy in AS. First, as PV+ neuron function appears critical for resilience to epileptogenesis, future work could aim to develop therapies to specifically modulate PV+ neuron activity or development. An allosteric modulator to enhance PV+ neuron firing has already shown preclinical efficacy in treating circuit dysfunction and sensory processing in a mouse model of Fragile X syndrome (Kourdougli et al., 2023), and would likely impact seizure phenotypes as well. Building on its role as an E3 ubiquitin ligase targeting protein substrates for degradation, future studies could also probe UBE3A substrates that are critical for PV+ neuron development and function (Mishra et al., 2009; Sell and Margolis, 2015; Krzeski et al., 2024). Identification of these molecules would inform strategies to modulate PV+ neuron function, which could drive additional treatments for epilepsies beyond AS. Finally, the sparse firing of DGCs is instrumental to these cells’ circuit contributions, allowing for pattern separation and the formation of place fields (Diamantaki et al., 2016; GoodSmith et al., 2017; Hainmueller and Bartos, 2020). Therefore, therapies designed to modulate DGC activity could improve seizure burden as well as other common and impactful comorbidities of epilepsy such as memory and mood disorders (Keezer et al., 2016; Hermann et al., 2021; Umschweif et al., 2021).

## Methods

### Animals

All procedures received approval from the Institutional Animal Care and Use Committee at the University of North Carolina at Chapel Hill and were conducted following the guidelines of the U.S. National Institutes of Health. Mice were kept on a 12:12 light/dark cycle (lights on at 7 A.M.) and housed in groups of 2-5 per cage with free access to food (PicoLab 5V5M chow) and water. Both male and female mice were used in roughly equal genotypic ratios. All experiments and post hoc data collection were carried out by experimenters blinded to genotype and treatment.

All mice were maintained on a C57BL/6J background for at least 10 generations. Mice with neuron type-specific deletion of *Ube3a* were generated by crossing male mice heterozygous for PV-T2A-Cre (JAX: 012358) (Madisen et al., 2012), SOM-IRES-Cre (JAX: 028864) (Taniguchi et al., 2011), VIP-IRES-Cre (JAX: 031628) (Taniguchi et al., 2011), or PV-IRES-Cre (JAX: 017320) (Hippenmeyer et al., 2005) with female mice heterozygous (paternal inheritance) or homozygous for the *Ube3a-Flox* allele (Judson et al., 2016). To generate mice with neuron type-specific reinstatement of *Ube3a*, male mice heterozygous for PV-T2A-Cre or Gad2-IRES-Cre (JAX: 028867) (Taniguchi et al., 2011) were crossed to females heterozygous (paternal inheritance) for the *Ube3a lox-STOP-lox* construct (Silva-Santos et al., 2015). Maternal *Ube3a*-deficient mice (*Ube3a^m-/p+^*; AS model mice) were produced by crossing female paternal *Ube3a*-deficient mice (JAX: 016590) to WT congenic C57BL/6J males.

To confirm cell type-specific deletion, we performed multiplex immunofluorescence for UBE3A and relevant lineage markers (PV, SOM, VIP) (Fig. S8). Quantification was conducted in QuPath using three sections per animal from four animals per condition. First, lineage marker-positive neurons were identified based on the marker and DAPI channels, after which their UBE3A status was assessed. Quantifying PV+ and SOM+ neurons in the DG and CA (Cornu Ammonis) regions showed that control mice maintained nearly complete UBE3A expression (99–100%). In deletion mutants, the Pvalb-T2A-Cre driver was highly effective, achieving 100% deletion in the DG and 97% in the CA. The SOM-IRES-Cre driver demonstrated comparable efficacy, with 96% and 88% deletion in the DG and CA, respectively. Conversely, the Pvalb-IRES-Cre line exhibited lower recombination efficiency, with 55% deletion in the DG and 78% in the CA. Although the scarcity of VIP+ interneurons prevented quantitative analysis, qualitative assessment confirmed UBE3A loss in most cells in VIP-IRES-Cre mice. Overall, these findings confirm the successful and specific removal of UBE3A in the targeted inhibitory cell populations.

### Flurothyl kindling

Mice aged 2-4 months were assessed for ictogenic and epileptogenic susceptibility using the flurothyl kindling and retest paradigm (Kadiyala et al. 2016). As previously described (Gu et al. 2019; Judson et al. 2021), mice were individually placed in a 2L airtight glass chamber and allowed to acclimate for 1 minute. After acclimation, 10% flurothyl (bis-2,2,2-trifluoroethyl ether), dissolved in 95% ethanol, was infused onto a filter paper disk (Whatman, Grade 1) suspended at the top of the chamber at 200 µL/min. Upon the start of a generalized seizure, characterized by limb clonus and loss of postural control, the chamber was opened to expose the mouse to fresh air. After behavioral seizures ceased, mice were placed in a clean recovery cage and then returned to their home cage. The chamber was cleaned with water and dried between mice, and cleaned with ethanol between cages.

Mice were subjected to flurothyl-induced seizures once daily for 8 consecutive days at a consistent time each day (induction phase), followed by a 28-day rest period during which no seizures were induced (incubation phase). On day 36, mice underwent one more flurothyl induction (retest). All seizure inductions were video recorded, and the latency to the first myoclonic jerk (myoclonic seizure threshold) and the onset of generalized seizure were recorded.

For electrophysiology experiments, *Ube3a^m-/p+^* mice and WT controls were kindled for 8 consecutive days or exposed to 8 consecutive days of 95% ethanol vapor for a comparable amount of time per day (sham kindled). Mice used for electrophysiology did not undergo a retest with flurothyl exposure. 32–65 days following flurothyl or sham kindling (experimental days 40–73), mice were selected for electrophysiology experiments, with one animal recording per day. The order of animal selection for electrophysiology experiments was assigned by an independent investigator and counterbalanced by genotype and treatment to minimize any effect of time since kindling.

### Immunohistochemistry

Mice were anesthetized with euthasol (100 mg/kg, i.p.) and transcardially perfused with PBS, followed by ice-cold 4% paraformaldehyde (PFA). Brains were post-fixed overnight in 4% PFA, cryoprotected in 30% sucrose, and sectioned coronally at 40 µm using a sliding microtome (Leica). Sections were stored at −20°C in cryopreservative solution (45% PBS, 30% ethylene glycol, 25% glycerol).

Free-floating sections were washed in PBS and blocked for 1 hour at room temperature in 5% normal goat serum containing 0.2% Triton X-100 (NGST). For VIP labeling, antigen retrieval was performed prior to blocking by incubating sections at 95°C for 30 minutes in 1 mM EDTA (pH 8.0). Sections were then incubated for 48 hours at 4°C with primary antibodies (Table S3). After PBS washes with 0.2% Triton-X, sections were incubated with fluorophore-conjugated secondary antibodies and counterstained with DAPI. Sections were mounted, air-dried, and coverslipped using Vectashield Plus (Vector Laboratories). Images were acquired using a Leica STELLARIS 8 FALCON or a Zeiss 710 confocal microscope, and a VS200 slide-scanning widefield microscope (Evident Scientific) with a 20x/0.8 objective.

### Image analysis

Image analysis was performed using QuPath (RRID: SCR_018257). WFA and GFAP labeling were quantified by measuring the mean fluorescence intensity of the corresponding channel in the molecular layer of the dentate gyrus. Images were acquired using a 20x/0.8 objective with a VS200 slide scanning widefield microscope (Evident Scientific). A background fluorescence was measured for WFA staining by taking the mean WFA fluorescence in the paraventricular nucleus of the thalamus, a region lacking noticeable perineuronal net expression, and this value was subtracted from the region of interest in the same slice. PV labeling was quantified by averaging fluorescence intensity across the entire hippocampus. Image acquisition for PV-IRES-Cre mice and controls was performed using a Zeiss 710 confocal microscope. Mean fluorescence values from all immunohistochemistry experiments represent averages from one hemisphere of 2-3 sections per mouse, and these values were normalized to controls for each staining run.

### Electrophysiology

#### Acute coronal slice preparation

32–65 days after flurothyl kindling (or sham kindling), mice were anesthetized with a lethal dose of Euthasol (100 mg/kg, intraperitoneal). After loss of toe pinch reflex, mice were transcardially perfused with ice-cold N-methyl-D-glucamine-based (NMDG) cutting solution bubbled with 95% O^2^ and 5% CO^2^ (in mM: 100 NMDG, 20 HEPES, 25 glucose, 30 NaHCO^3^, 5 sodium ascorbate, 2.5 KCl, 1.2 NaH^2^PO^4^-H^2^O, 2 thiourea, 3 sodium pyruvate, 5 N-acetyl-L-cysteine, 10 MgSO^4^-7H^2^O, 0.5 CaCl^2^-2H^2^O) (Ting et al., 2014). Brains were dissected and prepared in 300 µm-thick coronal slices in ice-cold NMDG cutting solution using a VT1200S vibratome (Leica). Slices recovered in NMDG solution at 34°C for 12 minutes in a submerged chamber bubbled with 95% O^2^ and 5% CO^2^, then transferred to a HEPES-buffered saline solution at room temperature (in mM: NaCl 92, 20 HEPES, 25 glucose, 30 NaHCO3, 5 sodium ascorbate, 2.5 KCl, 1.25 NaH^2^PO^4^-H^2^O, 2 thiourea, 3 sodium pyruvate, 5 N-acetyl-L-cysteine, 2 MgSO^4^-7H^2^O, 2 CaCl^2^-2H^2^O).

#### Whole-cell current clamp recordings

Slices containing dorsal hippocampus were placed in a submersion chamber perfused at 2-3 mL/min with carbogen-bubbled artificial cerebrospinal fluid (ACSF; in mM: 134 NaCl, 10 glucose, 25 NaHCO^3^, 5 myo-inositol, 0.4 sodium ascorbate, 3 KCl, 1.25 NaH^2^PO^4^-H^2^O, 2 sodium pyruvate, 1.3 MgSO^4^-7H^2^O, 2 CaCL^2^-2H^2^O) maintained at 30-32°C. Patch pipettes were pulled from fire polished borosilicate glass (1.5mm outer diameter, 0.86mm inner diameter) using a P2000 laser puller (Sutter Instruments), with open tip resistances of 4-7 MΩ. Pipettes were filled with internal solution containing the following (in mM): 130 potassium gluconate, 4 NaCl, 10 HEPES, 0.2 EGTA, 2 Mg-ATP, 0.3 Na-GTP, 10 tris phosphocreatine, pH adjusted to ∼7.3 with 2M KOH, and water was added if needed to adjust osmolarity to ∼290 mOSM.

Dentate granule cells were visually targeted using an Axio Examiner microscope (Zeiss) with infrared differential interference contrast imaging. Granule cells in the outer half of the suprapyramidal or infrapyramidal blade were chosen to avoid immature granule cells, and cells bordering the molecular layer were avoided to exclude semilunar granule cells. Cells with input resistance over 700 MΩ or resting membrane potential >-60 mV were excluded due to presumed immaturity. For successfully patched cells, we achieved membrane seal resistances >1 GΩ, and adjusted pipette capacitance compensation while in cell-attached configuration to minimize capacitive transients. Current clamp recordings were performed in whole-cell configuration using a Multiclamp 700B amplifier (Molecular Devices) with 100 kHz digitization and 20 kHz low-pass Bessel filtering. Series resistance was compensated using bridge balance mode. Series resistance was determined using a-10 mV voltage step and monitored throughout the recording. Cells were discarded if series resistance surpassed 25 MΩ.

Clampfit 11.4 was used for recording analysis. Input resistance was calculated with a-10 mV voltage step, using the steady state current before notable I^h^ current was observed. Resting membrane potential was recorded immediately after break-in. All membrane potentials are reported without correction for liquid junction potential.

To examine GC excitability, neurons were held at approximately −70 mV and were injected with 800 ms depolarizing current pulses (20 pA increments ranging up to 280pA, intertrial interval of at least 5 s). Action potentials crossing 0 mV were used for firing frequency analysis. Firing gain for each cell was quantified as the maximum increase in firing frequency over a 100 pA window of current steps. A separate protocol was used to calculate rheobase: beginning at −70 mV, cells were injected with 300 ms depolarizing current pulses in increments of 2 pA until they fired their first action potential. From this protocol, threshold voltage was defined as the point for which dVm/dt reached 10 mV/s.

#### Spontaneous inhibitory postsynaptic current recordings

To pharmacologically block excitatory transmission and isolate sIPSCs, we perfused sections at 30-32°C with ACSF containing 20 µM 6,7-dinitroquinoxaline-2,3-dione (DNQX) and 50 µM D-2-amino-5-phosphonopentanoic acid (D-APV). Patch pipettes were pulled from fire polished borosilicate glass (1.5mm outer diameter, 0.86mm inner diameter) using a P2000 laser puller (Sutter Instruments), with open tip resistances of 2.5-5 MΩ. Patch pipettes were filled with a high chloride Cs-based internal solution to amplify GABA-A receptor-mediated IPSCs (in mM: 65 cesium gluconate, 70 cesium chloride, 10 HEPES, 0.5 EGTA, 2 Mg-ATP, 0.3 Na-GTP, 10 disodium creatine phosphate, 5 QX-314 chloride). For all voltage-clamp experiments, series resistance was monitored throughout the recording, and cells were discarded if series resistance surpassed 20 MΩ.

To record sIPSCs, cells were held at −80 mV in voltage-clamp mode. Recordings were recorded at 20 kHz digitization and 6 kHz low-pass filtering, and were started at least 3 minutes after break-in to allow for sufficient diffusion of internal solution into the cell. Series resistance was not compensated for sIPSC recordings. To analyze sIPSC amplitude and frequency, 2 consecutive minutes of recordings were additionally low-pass filtered at 1.8 kHz, and the Clampfit template search function was used to extract individual events. In a subset of cells, 20µM gabazine was washed on the slice, which eliminated all inward current events, confirming that these were GABAergic sIPSCs.

#### Evoked inhibitory postsynaptic current recordings

eIPSCs were recorded under identical conditions to those described for sIPSCs using a high chloride Cs-based internal solution, −80 mV holding potential in voltage clamp mode, 20 kHz digitization, and 6 kHz low-pass filtering, and series resistance was compensated to 80%. IPSCs were electrically evoked using a concentric bipolar stimulating electrode (FHC, 125 µm tip diameter, Pt/Ir) placed in the middle of the granule cell layer ∼150 µm medial or lateral to the recorded granule cell. A stimulus duration of 200 µs was used. Stimulus intensity was controlled by a Digitimer DS4 bi-phasic stimulus isolation unit (Digitimer Limited). In a subset of cells, 20µM gabazine was applied to the slice, which eliminated all evoked currents.

Paired-pulse ratio was determined using inter-stimulus intervals of 25, 50, 100, and 200 ms, performed at each cell’s approximate half-max current. Synaptic facilitation was measured using a 20 Hz train of 10 pulses at each cell’s approximate half-max current with 10 seconds between each pulse train. One WT sham granule cell showed evoked current amplitudes over 200% of its baseline in the 20 Hz train protocol and was thus excluded from this analysis.

### RNA isolation and RT-qPCR

Brains were rapidly dissected from adult mice and flash frozen. RNA was isolated from whole hemispheres and extracted using RNeasy RNA extraction kit (Qiagen, 74104). cDNA was prepared using qScript cDNA SuperMix (QuantaBio, 95048-100) from 1 µg input RNA. PowerUp SYBR Mastermix (Applied Biosytems, A25742) was used for RTqPCR on the QuantStudio5 (Applied Biosystems). Relative *Pvalb* expression was analyzed using the 2−ddCT method, normalized to *Actb.* Primers used: pvalb_mus1_F: GGCCTGAAGAAAAAGAACCCGG, pvalb_mus1_R: GACAAGTCTCTGGCATCTGAGG, ACTB_F: GGCACCACACCTTCTACAATG, ACTB_R: GGGGTGTTGAAGGTCTCAAAC.

### Western blotting

Brains were rapidly dissected from adult mice, flash frozen, and stored at −80 °C until lysing. Tissues were lysed in RIPA buffer (50 mM Tris-HCl, 150 mM NaCl, 1% NP-40, 0.5% Sodium Deoxycholate), 0.01% protease inhibitor (P8340 Millipore Sigma), and 0.5% SDS. After homogenization with a Tissue Tearor (Model 985-370), samples were centrifuged for 10 min at 21,000 g, 4°C. Protein concentration was measured from the supernatant using the Pierce BCA kit (Thermo Scientific). 20 µg of each sample was loaded into a 10% SDS polyacrylamide gel (Mini-PROTEAN TGX precast gel) submerged in Tris-Glycine/SDS buffer (25 mM Tris-base, 192 mM glycine, 0.1% SDS). Samples were transferred onto a 0.2 µm nitrocellulose membrane (Bio-Rad) ran for 60 min at 100 V in ice cold transfer buffer (25 mM Tris-base, 192 mM glycine, 20% MeOH). Membranes were incubated in Intercept blocking buffer (Li-Cor) and subsequently incubated overnight with primary antibodies Rabbit anti-Parvalbumin (1:1000, Swant, PV27a) or Mouse anti-GAPDH (1:5000, Sigma, MAB374) at 4°C. The next day, membranes were washed for 30 min with PBS/0.1% Tween-20 and incubated in HRP-conjugated secondary antibodies (1:5000 Invitrogen, anti-mouse 31430 or anti-rabbit 31460) for 1hr at room temperature, followed by 30 min washing with PBS/0.1% Tween-20). The membranes were imaged using Clarity Western ECL (Bio-Rad) substrate on an Amersham Imager (AI600, GE Life Sciences). Uncropped western blot images presented in Fig. S9.

### Experimental Design and Statistical Analysis

Two-way repeated measures ANOVAs were used for flurothyl kindling seizure behavior analysis. One-way ANOVAs were used for hippocampal histology experiments. Three-way ANOVAs with were used for electrophysiological measures across multiple intensities (frequency-current curves, evoked inhibitory input-output curves, paired-pulse ratio, 20 Hz pulse train), and two-way ANOVAs were used for cellular properties with one value per cell (input resistance, rheobase, resting membrane potential, etc.). For all ANOVAs with significant main effects or interactions, Šídák’s multiple comparisons *post hoc* tests were used when comparing means of two experimental groups, and Tukey’s *post hoc* tests were used when comparing means of three or more groups. For comparisons between two group means, two-tailed unpaired t-tests or two-tailed Welch’s tests were used, depending on whether the groups had significantly different variances on F test. Statistical tests used for each experiment are included in the corresponding figure legends. Data are presented as means ± SEM unless otherwise noted. A *P*-value less than 0.05 was considered statistically significant. Sample sizes needed to achieve statistical significance (p<0.05) were estimated using power analyses and expected effect size based on previously published and preliminary data. Graphpad Prism 10.3.1 software was used for all statistical analyses.

## Acknowledgements

This work was supported by the Simons Foundation Autism Research Initiative (SFARI 702556), NIH R01NS129914, NIH R01NS131615, NIH R01NS145518, and NIH F30HD111296. We thank Wendy Salmon and the UNC Hooker Imaging Core Facility, supported in part by P30 CA016086 Cancer Center Core Support Grant to the UNC Lineberger Comprehensive Cancer Center. Microscopy was performed at the UNC Neuroscience Microscopy Core (RRID:SCR_019060), supported, in part, by funding from the NIH-NICHD Intellectual and Developmental Disabilities Research Center Support Grant P50 HD103573.

---

**Supplemental Figure 1: Data from Figure 1 with control groups disaggregated.** Latency to (**A**) myoclonic seizure and (**B**) generalized seizure in WT mice and *Ube3a^mFLOX/p+^* mice. Two-way RM ANOVA with Šídák’s post hoc tests. (**C,D**) Seizure latencies for PV-T2A-Cre:*Ube3a^mFLOX/p+^* mice and controls. Two-way RM ANOVA with Tukey’s post hoc tests. For clarity, only statistically significant post hoc comparisons between WT and Cre+ controls are shown. (**E,F**) Seizure latencies for SOM-Cre:*Ube3a^mFLOX/p+^* mice and controls. Data presented as means ± SEM. *P < 0.05, **P < 0.01, ***P<0.001.

**Supplemental Figure 2: PV-IRES-Cre mice show resistance to seizure kindling and decreased expression of parvalbumin.** Latency to (**A**) myoclonic seizure and (**B**) generalized seizure in WT, *Ube3a^mFLOX/p+^*, PV-IRES-Cre, and PV-IRES-Cre:*Ube3a^mFLOX/p+^* mice. Two-way RM ANOVA with Tukey’s post hoc tests. Asterisks represent difference compared to WT control, color coded by comparison group. (**C**) Representative images of parvalbumin staining in hippocampus of WT and PV-IRES-Cre mice. (**D**) Quantification of mean fluorescence of parvalbumin staining across hippocampus of flurothyl kindled mice. One-way ANOVA with Tukey’s post hoc tests. (**E**) Quantification of mean fluorescence of parvalbumin staining in hippocampus of non-kindled WT and PV-IRES-Cre mice. Unpaired t-test. Data presented as means ± SEM. *P < 0.05, **P < 0.01, ***P<0.001.

**Supplemental Figure 3: PV-IRES-Cre mice exhibit decreased parvalbumin transcript and protein levels.** (**A**) RT-qPCR for *Pvalb* levels in hemispheres of WT and hemizygous PV-IRES-Cre mice. Unpaired t-test. (**B**) Western blot for parvalbumin levels in hemispheres of WT, hemizygous, and homozygous PV-IRES-Cre. One-way ANOVA with Tukey’s post hoc tests. Data presented as means ± SEM. *P < 0.05, **P < 0.01, ***P<0.001.

**Supplemental Figure 4: PV-T2A-Cre mice show decreased parvalbumin expression.** (**A**) RT-qPCR for *Pvalb* levels in hemispheres of WT, hemizygous and homozygous PV-T2A-Cre mice. Brown-Forsythe Test with Dunnett’s T3 multiple comparisons tests. (**B**) Quantification of mean PV fluorescence in hippocampus of WT and PV-T2A-Cre mice. Unpaired t-test. Data presented as means ± SEM. *P < 0.05, **P < 0.01, ***P<0.001.

**Supplemental Figure 5: Data from Figure 2 with control groups disaggregated.** (**A**) Myoclonic and (**B**) generalized seizure latencies in Gad2-Cre:*Ube3a^mSTOP/p+^* mice and controls. (**C**) Seizure mortality in Gad2-Cre:*Ube3a^mSTOP/p+^* mice and controls. (**D**) Myoclonic and (**E**) generalized seizure latencies for PV-T2A-Cre:*Ube3a^mSTOP/p+^* mice and controls. (**F**) Seizure mortality in PV-T2A-Cre:*Ube3a^mSTOP/p+^* mice and controls. (**A,B,D,E**) Two-way RM ANOVA with Tukey’s post hoc tests. (**C,F**) Log-rank (Mantel-Cox) test. For clarity, only statistically significant post hoc comparisons between WT and Cre+ controls are shown. Data presented as means ± SEM. *P < 0.05, **P < 0.01, ***P<0.001.

**Supplemental Figure 6: Reactive astrocytosis in the dentate gyrus correlates with seizure susceptibility.** (**A**) Representative images of GFAP immunofluorescence in the hippocampus of WT, *Ube3a^mSTOP/p+^*, and PV-T2A-Cre:*Ube3a^mSTOP/p+^* mice (Scale bar: 250 µm). (**B**) Mean GFAP fluorescence intensity in DG molecular layer, normalized to controls, in flurothyl kindled Gad2-Cre:*Ube3a^mSTOP/p+^* mice, (**C**) PV-T2A-Cre:*Ube3a^mFLOX/p+^* mice, and (**D**) PV-T2A-Cre:*Ube3a^mSTOP/p+^* mice. Green data points in (**D**) correspond to example images in (**A**). (**E**) Correlation analysis of DG GFAP fluorescence and myoclonic seizure threshold of controls and *Ube3a^mSTOP/p+^* mice. (**F**) Correlation analysis of DG GFAP fluorescence and generalized seizure threshold. (**B,D**) One-way ANOVA with Tukey’s post hoc tests. (**C**) Unpaired t-test. (**E,F**) Simple linear regression, with * indicating significantly non-zero slope. Data presented as means ± SEM. *P < 0.05, **P < 0.01, ***P<0.001.

**Supplemental Figure 7: Representative traces of ‘doublet’ firing dentate granule cells.** (**A-C**) Representative current-clamp traces from three doublet firing dentate granule cells from kindled AS mice upon current injection of (**i**) 120 pA, and (**ii**) 160 pA. (**D**) Current-clamp trace from sham-kindled AS mouse demonstrating single doublet firing on one current injection. (**i**) 260 pA, (**ii**) 280 pA. Scale bars: (**i, ii**) 25 mV, 100 ms, (**iii**) 5 mV, 5 ms.

**Supplemental Figure 8: Cell type-specific removal of UBE3A in hippocampal interneuron subtypes.** Representative immunofluorescence images showing UBE3A expression (red) in (**A**) PV+, (**B**) SOM+, and (**C**) VIP+ interneurons (green) in the hippocampal CA1 region of control and Cre-mediated deletion mice. In control mice, cell type-specific markers colocalize with UBE3A (dashed outlines). In deletion mice, targeted interneuron subtypes express their respective markers but lack UBE3A expression (dashed outlines indicate marker-positive, UBE3A-negative cells), while surrounding non-targeted cells retain UBE3A expression. PV-T2A-Cre and SOM-IRES-Cre mice show robust deletion, with the majority of labeled interneurons lacking UBE3A. VIP-IRES-Cre mice also demonstrate loss of UBE3A in VIP+ cells, though these cells are sparse in the hippocampus. SO, stratum oriens; SP, stratum pyramidale; SR, stratum radiatum. Scale bars, 50 μm.

**Supplemental Figure 9: Uncropped images of western blots** (**A**) Uncropped image of western blot for parvalbumin of WT, PV-IRES-Cre hemizygous, and PV-IRES-Cre homozygous lysates of brain hemispheres. (**B**) Uncropped image of western blot for GAPDH in same samples.

