## Supplemental Figures and Tables for "Parvalbumin interneurons and dentate gyrus homeostatic dysregulation shape epileptogenesis in Angelman syndrome model mice"

Fig. S1

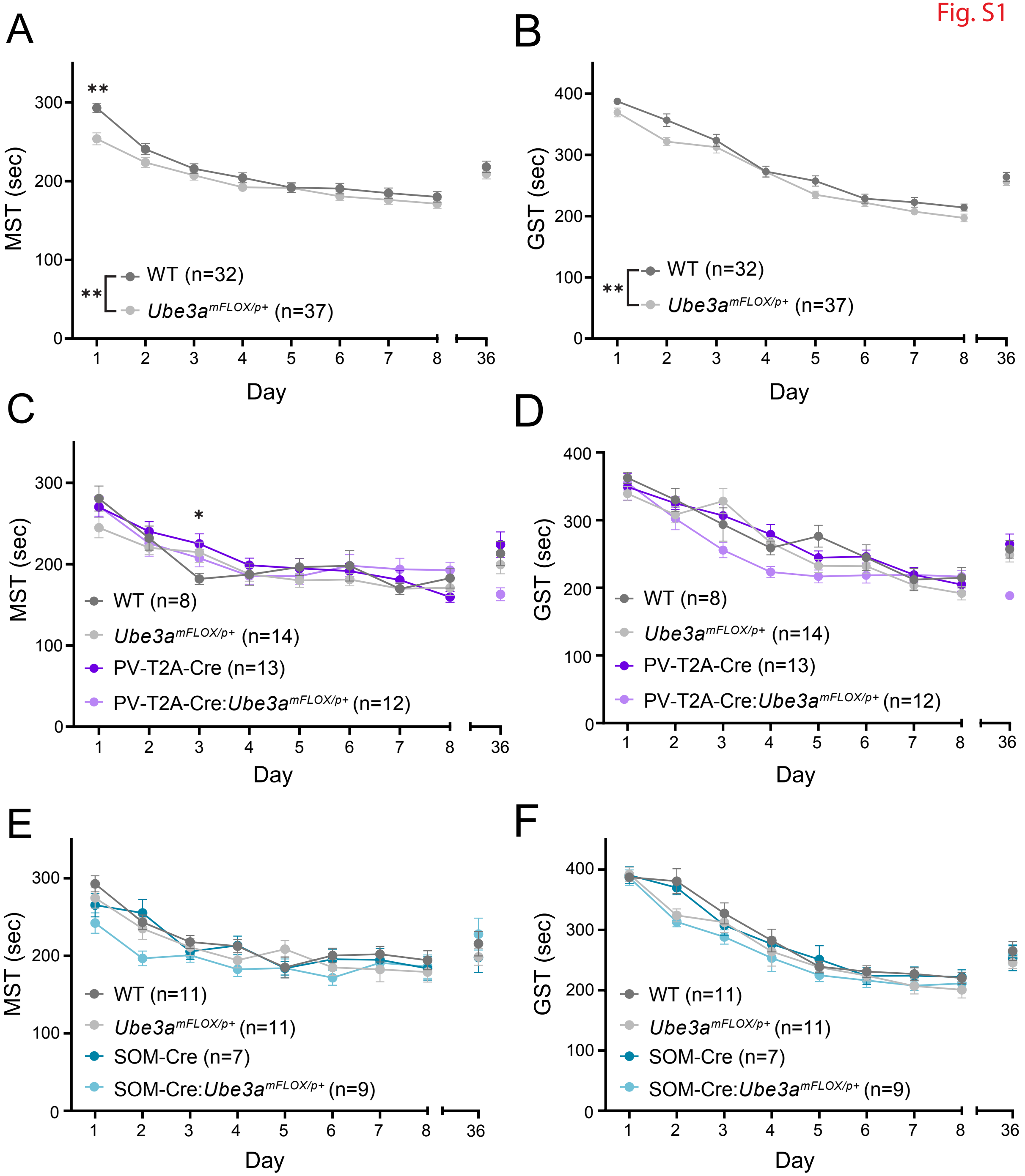
A
MST (sec)
Day
WT (n=32)
Ube3a^mFLOX/p+ (n=37)
**
B
GST (sec)
Day
WT (n=32)
Ube3a^mFLOX/p+ (n=37)
**
C
MST (sec)
Day
*
WT (n=8)
Ube3a^mFLOX/p+ (n=14)
PV-T2A-Cre (n=13)
PV-T2A-Cre:Ube3a^mFLOX/p+ (n=12)
D
GST (sec)
Day
WT (n=8)
Ube3a^mFLOX/p+ (n=14)
PV-T2A-Cre (n=13)
PV-T2A-Cre:Ube3a^mFLOX/p+ (n=12)
E
MST (sec)
Day
WT (n=11)
Ube3a^mFLOX/p+ (n=11)
SOM-Cre (n=7)
SOM-Cre:Ube3a^mFLOX/p+ (n=9)
F
GST (sec)
Day
WT (n=11)
Ube3a^mFLOX/p+ (n=11)
SOM-Cre (n=7)
SOM-Cre:Ube3a^mFLOX/p+ (n=9)

Fig. S2

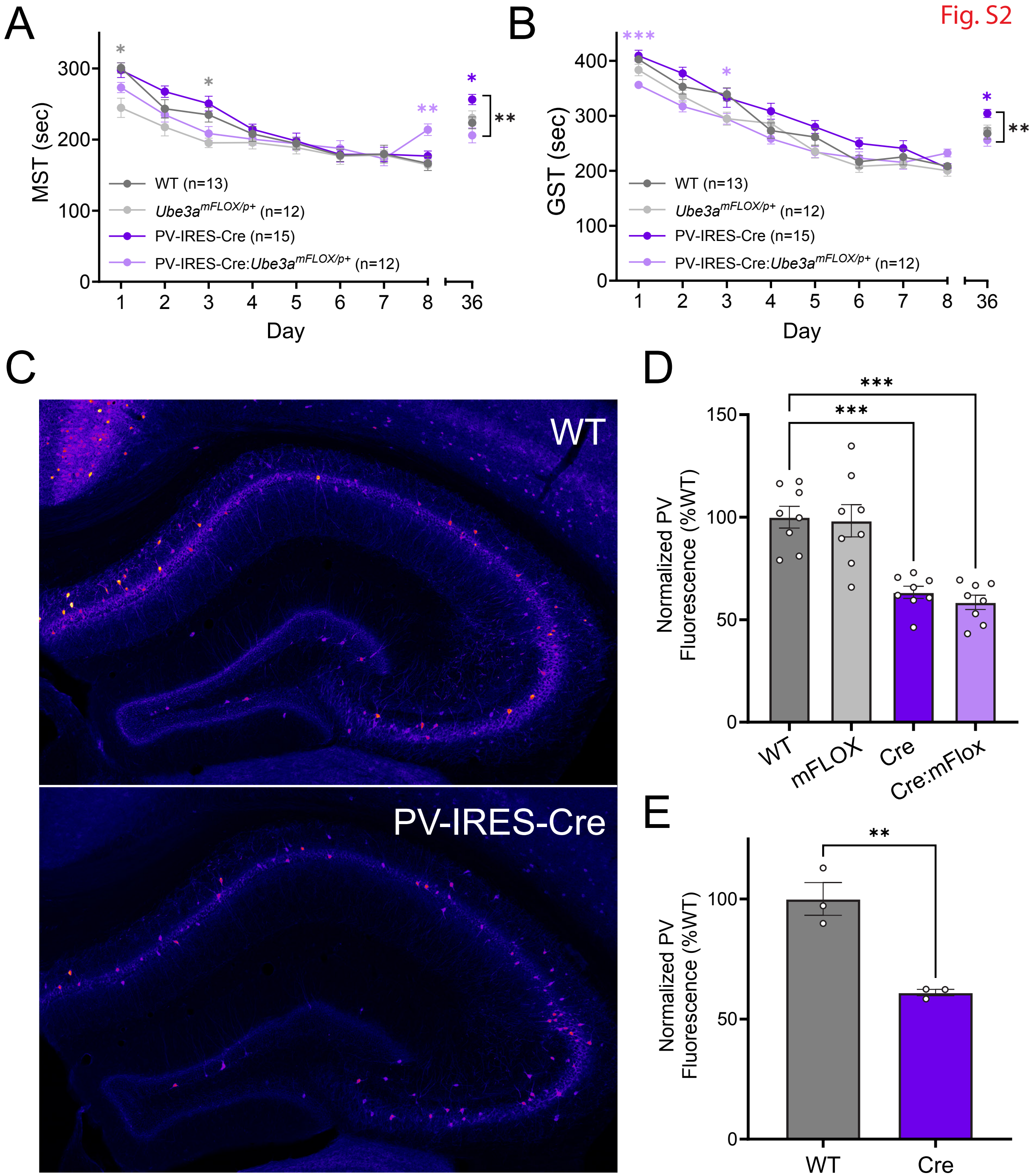

A
MST (sec)
Day
WT (n=13)
Ube3a^mFLOX/p+ (n=12)
PV-IRES-Cre (n=15)
PV-IRES-Cre:Ube3a^mFLOX/p+ (n=12)
B
GST (sec)
C
WT
PV-IRES-Cre
D
Normalized PV Fluorescence (%WT)
WT
mFLOX
Cre
Cre:mFlox
E
Normalized PV Fluorescence (%WT)
WT
Cre

### PV-IRES-Cre

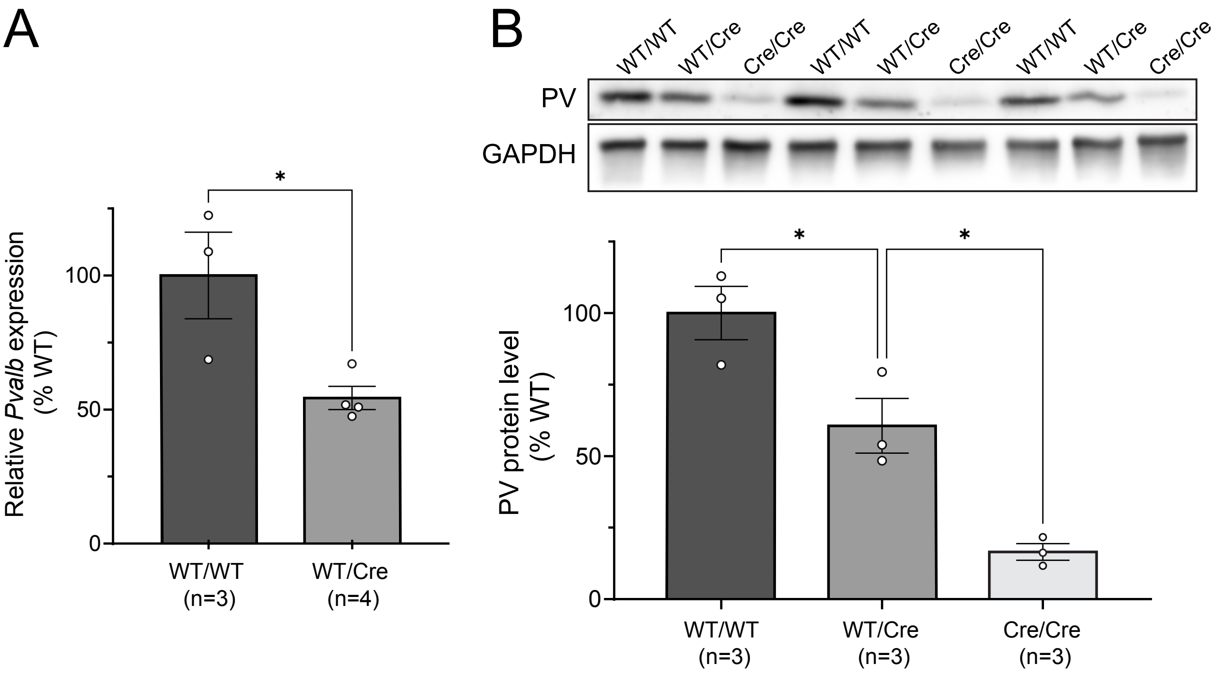
A
Relative *Pvalb* expression (% WT)
0
50
100
*
WT/WT (n=3)
WT/Cre (n=4)
B
WT/WT
WT/Cre
Cre/Cre
WT/WT
WT/Cre
Cre/Cre
WT/WT
WT/Cre
Cre/Cre
PV
GAPDH
PV protein level (% WT)
0
50
100
*
*
WT/WT (n=3)
WT/Cre (n=3)
Cre/Cre (n=3)

Fig. S4

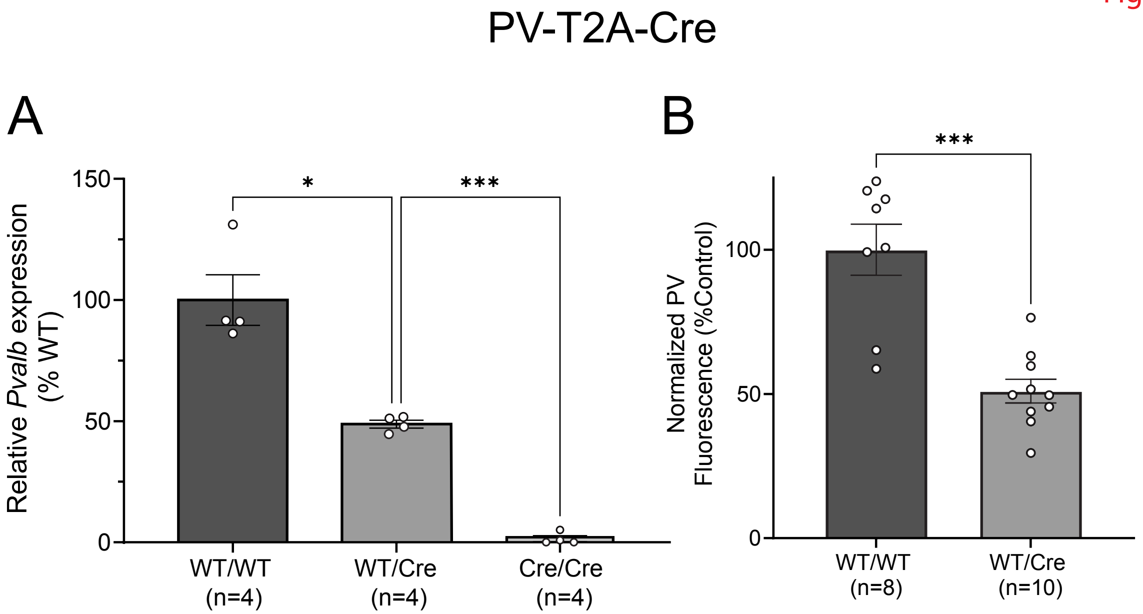
PV-T2A-Cre
A
Relative *Pvalb* expression (% WT)
150
100
50
0
*
***
WT/WT (n=4)
WT/Cre (n=4)
Cre/Cre (n=4)
B
Normalized PV Fluorescence (%Control)
100
50
0
***
WT/WT (n=8)
WT/Cre (n=10)

Fig. S5

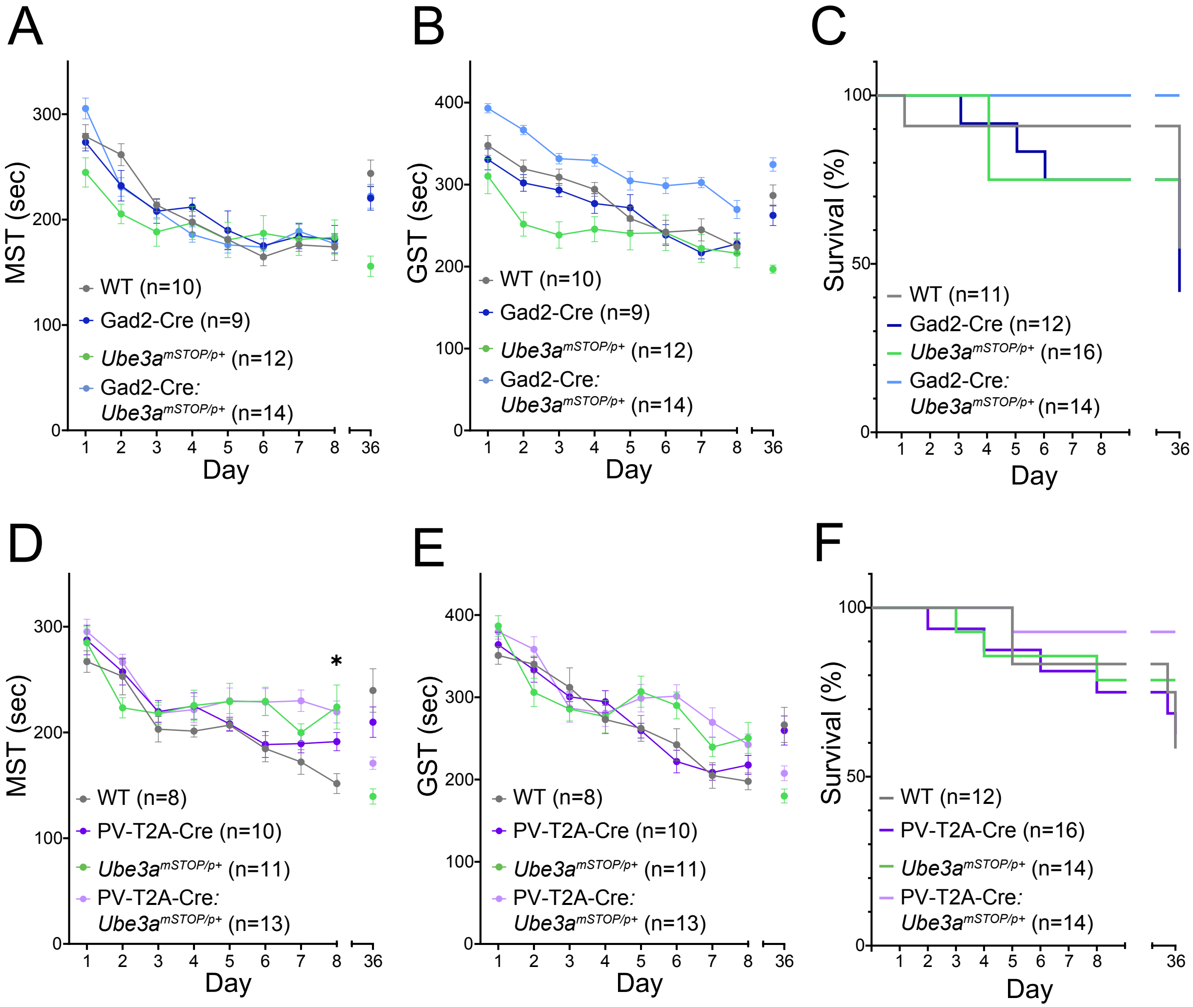
A
MST (sec)
Day
WT (n=10)
Gad2-Cre (n=9)
Ube3a^mSTOP/p+ (n=12)
Gad2-Cre: Ube3a^mSTOP/p+ (n=14)
B
GST (sec)
Day
WT (n=10)
Gad2-Cre (n=9)
Ube3a^mSTOP/p+ (n=12)
Gad2-Cre: Ube3a^mSTOP/p+ (n=14)
C
Survival (%)
Day
WT (n=11)
Gad2-Cre (n=12)
Ube3a^mSTOP/p+ (n=16)
Gad2-Cre: Ube3a^mSTOP/p+ (n=14)
D
MST (sec)
Day
WT (n=8)
PV-T2A-Cre (n=10)
Ube3a^mSTOP/p+ (n=11)
PV-T2A-Cre: Ube3a^mSTOP/p+ (n=13)
E
GST (sec)
Day
WT (n=8)
PV-T2A-Cre (n=10)
Ube3a^mSTOP/p+ (n=11)
PV-T2A-Cre: Ube3a^mSTOP/p+ (n=13)
F
Survival (%)
Day
WT (n=12)
PV-T2A-Cre (n=16)
Ube3a^mSTOP/p+ (n=14)
PV-T2A-Cre: Ube3a^mSTOP/p+ (n=14)

Fig. S6

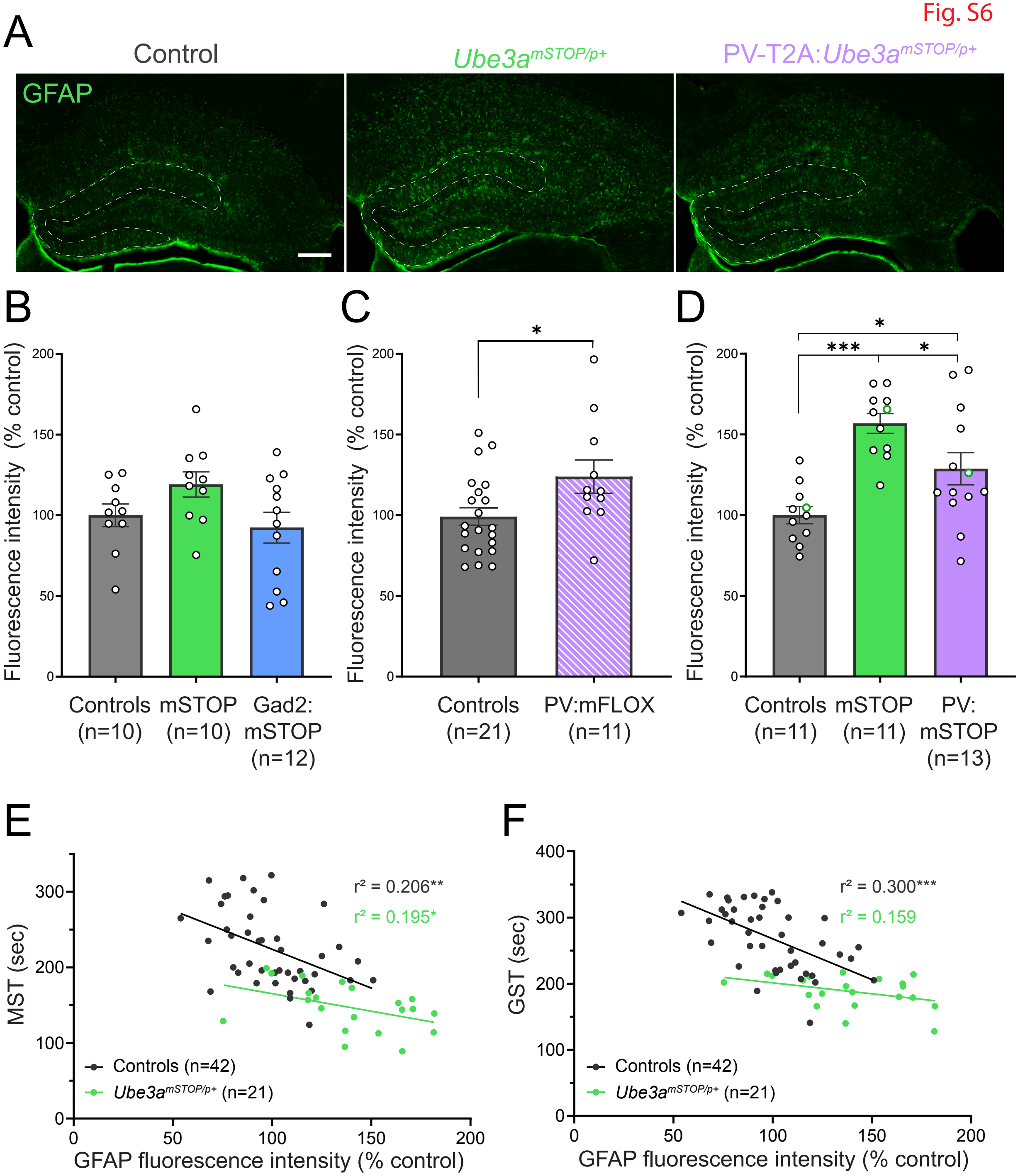

A
Control
Ube3a mSTOP/p+
PV-T2A:Ube3a mSTOP/p+
GFAP
B
Fluorescence intensity (% control)
Controls (n=10)
mSTOP (n=10)
Gad2: mSTOP (n=12)
C
Fluorescence intensity (% control)
Controls (n=21)
PV:mFLOX (n=11)
D
Fluorescence intensity (% control)
Controls (n=11)
mSTOP (n=11)
PV: mSTOP (n=13)
E
MST (sec)
r² = 0.206**
r² = 0.195*
Controls (n=42)
Ube3a mSTOP/p+ (n=21)
GFAP fluorescence intensity (% control)
F
GST (sec)
r² = 0.300***
r² = 0.159
Controls (n=42)
Ube3a mSTOP/p+ (n=21)
GFAP fluorescence intensity (% control)

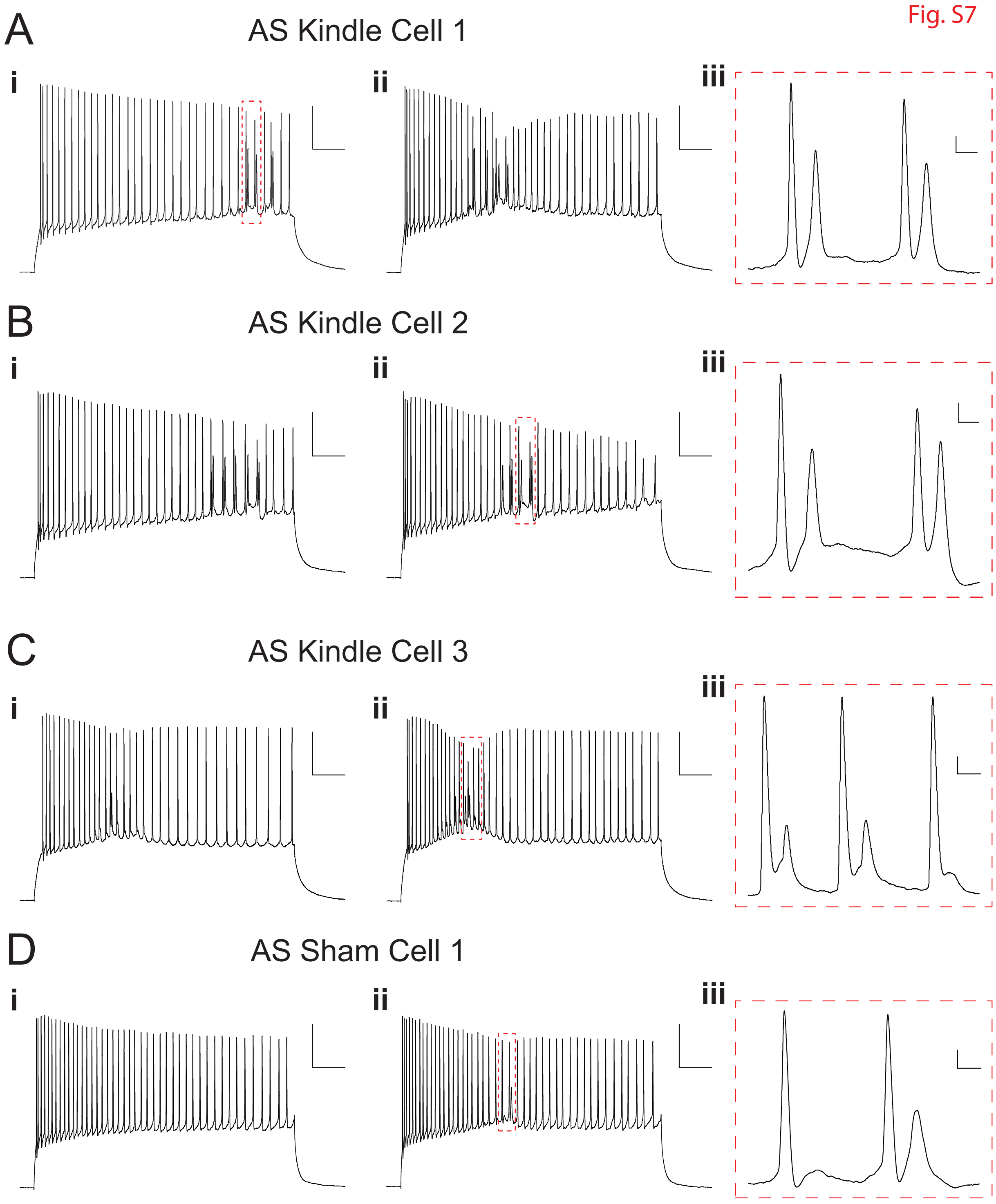
Fig. S7
A
AS Kindle Cell 1
i
ii
iii
B
AS Kindle Cell 2
i
ii
iii
C
AS Kindle Cell 3
i
ii
iii
D
AS Sham Cell 1
i
ii
iii

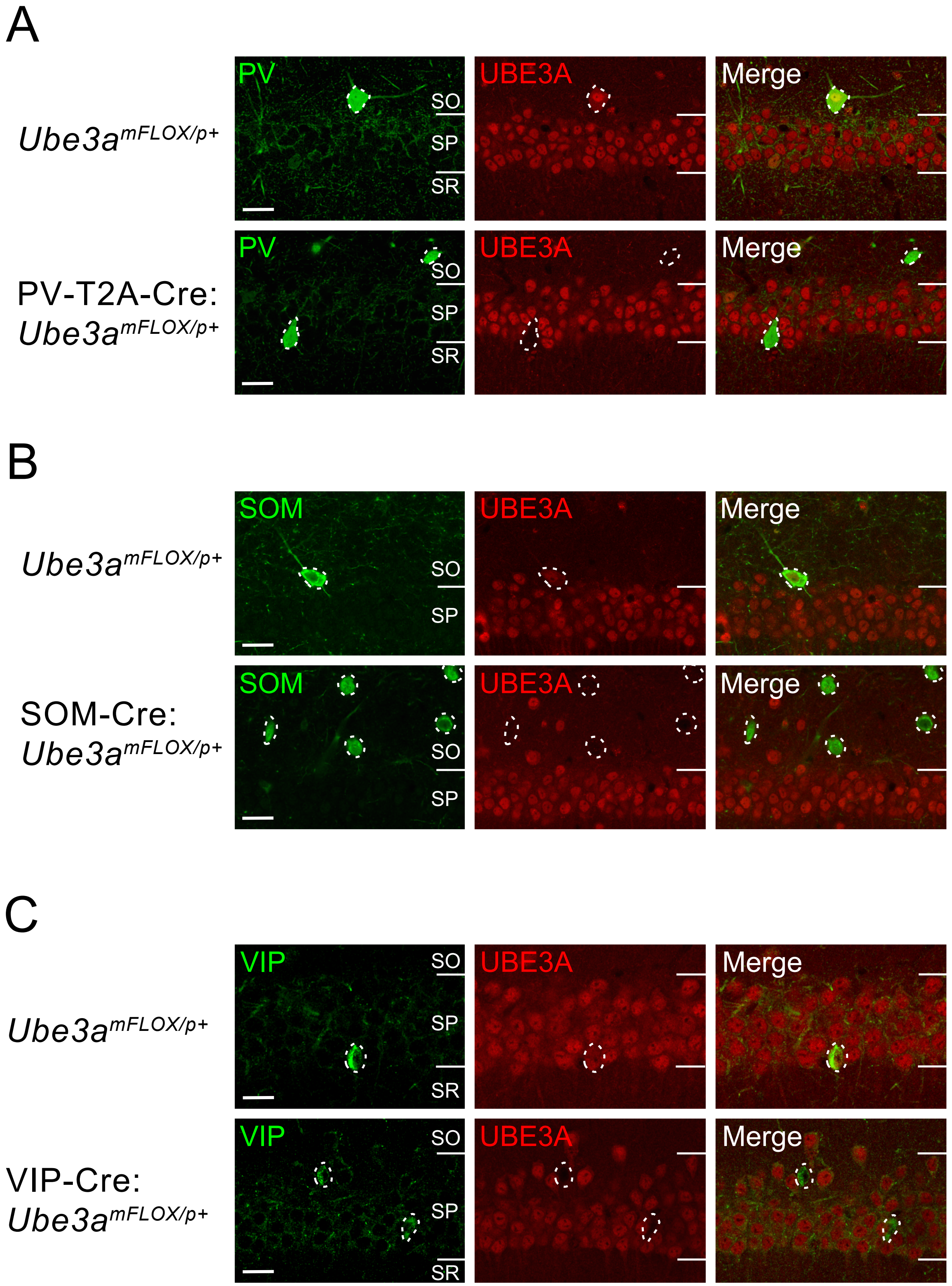
A
$Ube3a^{mFLOX/p+}$
PV-T2A-Cre:
$Ube3a^{mFLOX/p+}$
PV
UBE3A
Merge
SO
SP
SR
B
$Ube3a^{mFLOX/p+}$
SOM-Cre:
$Ube3a^{mFLOX/p+}$
SOM
UBE3A
Merge
SO
SP
C
$Ube3a^{mFLOX/p+}$
VIP-Cre:
$Ube3a^{mFLOX/p+}$
VIP
UBE3A
Merge
SO
SP
SR

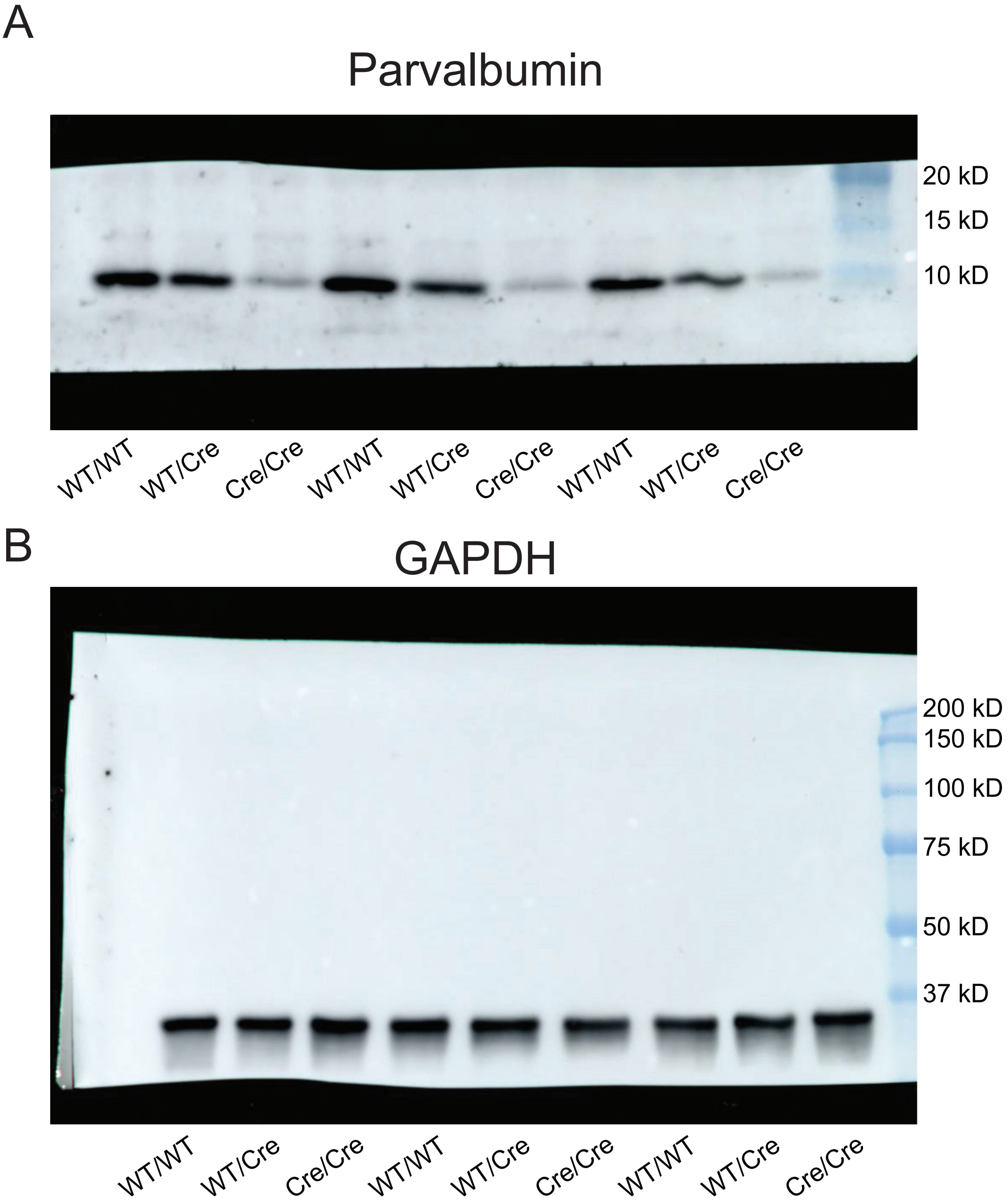
A
Parvalbumin
20 kD
15 kD
10 kD
WT/WT
WT/Cre
Cre/Cre
WT/WT
WT/Cre
Cre/Cre
WT/WT
WT/Cre
Cre/Cre
B
GAPDH
200 kD
150 kD
100 kD
75 kD
50 kD
37 kD
WT/WT
WT/Cre
Cre/Cre
WT/WT
WT/Cre
Cre/Cre
WT/WT
WT/Cre
Cre/Cre

**Supplemental Table 1. Intrinsic properties of dentate granule cells**

| Property | WT Sham (n=18) | WT Kindle (n=25) | AS Sham (n=18) | AS Kindle (n=25) | Genotype | Condition | Interaction |
|---|---|---|---|---|---|---|---|
| Input resistance (MΩ) | 315 ± 22 | 360 ± 20 | 297 ± 18 | 395 ± 17 | 0.6625 | *0.0004 | 0.1748 |
| Resting membrane potential (mV) | -78.2 ± 1.0 | -78.1 ± 0.99 | -76.5 ± 1.3 | -78.2 ± 0.75 | 0.4128 | 0.4547 | 0.3605 |
| Rheobase (pA) | 60 ± 6.1 | 55 ± 3.7 | 67 ± 7.5 | 48 ± 4.5 | 0.9897 | *0.0257 | 0.176 |
| AP threshold (mV) | -40.3 ± 1.0 | -39.1 ± 1.1 | -38.0 ± 1.3 | -38.2 ± 1.3 | 0.2118 | 0.7127 | 0.5814 |
| Firing gain (Hz/nA) | 229.2 ± 14.4 | 232.5 ± 14.4 | 211.1 ± 19.5 | 297.0 ± 25.2 | 0.2483 | *0.0282 | *0.0419 |
| Max firing frequency (Hz) | 41.9 ± 2.8 | 39.2 ± 2.5 | 35.1 ± 3.4 | 41.5 ± 3.3 | 0.4620 | 0.5546 | 0.1391 |
| Spike Frequency Adaptation Ratio | 0.392 ± 0.063 | 0.398 ± 0.042 | 0.391 ± 0.063 | 0.571 ± 0.062 | 0.1440 | 0.1156 | 0.1412 |

Data presented as mean ± SEM. Two-way ANOVA for effect of genotype and seizure kindling. *$P < 0.05$

**Supplemental Table 2. Intrinsic properties of 'doublet' firing dentate granule cells in kindled AS mice**

| Property | Non-doublet firing (n=19) | Doublet firing (n=6) | Unpaired t test / Welch's t test |
|---|---|---|---|
| Input resistance (MΩ) | 387 ± 19 | 422 ± 40 | 0.3884 |
| Resting membrane potential (mV) | -77.9 ± 0.95 | -79.0 ± 0.82 | 0.5511 |
| Rheobase (pA) | 52 ± 5.5 | 33 ± 2.4 | *#0.0039 |
| AP threshold (mV) | -36.4 ± 1.5 | -44.0 ± 1.2 | *0.0130 |
| Firing gain (Hz/nA) | 267.8 ± 29.2 | 389.6 ± 27.7 | *0.0361 |
| Max firing frequency (Hz) | 37.8 ± 3.8 | 53.1 ± 3.6 | *0.0432 |
| Spike Frequency Adaptation Ratio | 0.549 ± 0.075 | 0.638 ± 0.11 | 0.5550 |

Data presented as mean ± SEM. Unpaired t test or Welch's t test comparing doublet and non-doublet firing cells. *P < 0.05
### Welch's t test used for comparison of groups with unequal variance.

#### Supplemental Table 3. Antibodies used

| | | | Primary Antibody | Secondary Antibody |
|---|---|---|---|---|
| **Microscopy** | | | | |
| Figure S8 | **Parvalbumin** | Antibody | anti-Parvalbumin | anti-guinea pig, Alexa Fluor 488 |
| | | Host, type | Guinea pig, monoclonal (Gp58E1) | Donkey, polyclonal |
| | | Dilution | 1/2,000 | 1/400 |
| | | Company, #Cat. | Synaptic Systems, 195 308 | Jackson ImmunoResearch Labs, 706-545-148 |
| | | Reference | RRID: AB_2927389 | RRID: AB_2340472 |
| | **Somatostatin** | Antibody | anti-Somatostatin-28 | anti-rat, Alexa Fluor Plus 647 |
| | | Host | Rat, monoclonal (SY-160F7) | Donkey, polyclonal |
| | | Dilution | 1/1,000 | 1/400 |
| | | Company, #Cat. | Synaptic Systems, 366 017 | Thermo Fisher Scientific, A48272TR |
| | | Reference | RRID: AB_3083022 | RRID: AB_2896338 |
| | **VIP** | Antibody | anti-Vasoactive intestinal peptide | anti-rabbit, Alexa Fluor Plus 647 |
| | | Host, type | Rabbit, monoclonal (EPR23288-43) | Donkey, polyclonal |
| | | Dilution | 1/2,000 | 1/400 |
| | | Company, #Cat. | Abcam, ab272726 | Thermo Fisher Scientific, A32795TR |
| | | Reference | RRID:AB_3718106 | RRID:AB_2866496 |
| | **UBE3A** | Antibody | anti-UBE3A | anti-mouse IgG, Alexa Fluor Plus 594 |
| | | Host, type | Mouse, monoclonal | Donkey, polyclonal |
| | | Dilution | 1/1,000 | 1/400 |
| | | Company, #Cat. | Sigma-Aldrich, SAB1404508 | Thermo Fisher Scientific, A32744 |
| | | Reference | RRID:AB_10740376 | RRID:AB_2762826 |
| Figure 3, S6 | **WFA** | Antibody<br>Host, type | Biotinylated *Wisteria floribunda* agglutinin | Streptavidin, Alexa Fluor 568 |
| | | Dilution | 1/1,000 | 1/500 |
| | | Company, #Cat. | Sigma, L1516 | Thermo Fisher Scientific, S11226 |
| | **GFAP** | Antibody | anti-Glial fibrillary acidic protein | anti-rabbit, Alexa Fluor 488 |
| | | Host, type | Rabbit, polyclonal | Goat, polyclonal |
| | | Dilution | 1/1,000 | 1/500 |
| | | Company, #Cat. | DAKO, Z0334 | Thermo Fisher Scientific, A-11008 |
| | | Reference | RRID: AB_10013382 | RRID:AB_143165 |
| Fig. S2, S3, S4 | **Parvalbumin** | Antibody | anti-Parvalbumin | anti-mouse IgG1, Alexa Fluor 647 |
| | | Host, type | Mouse, monoclonal | Goat, polyclonal |
| | | Dilution | 1/2,000 | 1/500 |
| | | Company, #Cat. | Swant, PV235 | Thermo Fisher Scientific, A-21240 |
| | | Reference | RRID:AB_3698492 | RRID:AB_2535809 |
| **Western Blotting** | | | | |
| Figure S3 | **Parvalbumin** | Antibody | anti-Parvalbumin | anti-rabbit, HRP-conjugated |
| | | Host, type | Rabbit, polyclonal | Goat, polyclonal |
| | | Dilution | 1/1,000 | 1/5,000 |
| | | Company, #Cat. | Swant, PV27a | Thermo Fisher Scientific, 31460 |
| | | Reference | RRID:AB_3741048 | RRID:AB_228341 |
| | **GAPDH** | Antibody | anti-Glyceraldehyde-3-PDH | anti-mouse, HRP-conjugated |
| | | Host, type | Mouse, monoclonal | Goat, polyclonal |
| | | Dilution | 1/5,000 | 1/5,000 |
| | | Company, #Cat. | Sigma, MAB374 | Thermo Fisher Scientific, 31430 |
| | | Reference | RRID:AB_2107445 | RRID:AB_228307 |
